# Multi-omics analysis of TNBC organoids identifies phosphorylation of the membrane trafficking machinery as key event associated with FER-mediated invasion

**DOI:** 10.64898/2025.12.18.695070

**Authors:** Carolien Elshout, Gonçalo Mota, Joana Cunha, Sara Largher, Paula Sobrevals Alcaraz, Andreia Silva, Bruno Cavadas, Peter D. Haughton, Tuyet Anh Nguyen, Robert M. Van Es, Bruno S. Monteiro, Ana Rita Araújo, Ana Marques, Marta T. Pinto, Maria Peixoto, Sven van Kempen, Kevin van der Graaf, Anne-Marie Fortier, Morag Park, Paul J. van Diest, Antoine A. Khalil, Rui Fernandes, Harmjan R. Vos, Patrick W. B. Derksen, Florence Janody, Sandra Tavares

## Abstract

Triple Negative Breast Cancer (TNBC) is characterised by unfavourable outcome due to the combination of its metastatic propensity, chemo-refractory behaviour and the lack of effective targeted interventions. Expression of the feline sarcoma-related (FER) kinase constitutes an independent prognostic factor that correlates with poor patient survival. FER promotes invasive behaviour in TNBC cells by regulating endosomal sorting and recycling (ESR) of adhesion proteins. Yet, the ESR machinery supporting invasion in TNBC, particularly within 3D environments, remains poorly understood. Here, we used FER-expressing TNBC patient-derived xenograft organoids (PDXOs) and MDA-MB-231 cells to identify the membrane trafficking machinery promoting invasion. Using a combination of proteomics, phospho-proteomics, and single cell RNA-sequencing, we show that the invasion of FER-expressing PDXO cells in collagen-I is mainly associated with the differential phosphorylation of membrane trafficking regulators, including SEC16A and a marked increase in Rab4-positive tubules. SEC16A depletion impairs cell invasion and reduces the number of focal adhesions and Rab4-positive tubules. Importantly, FER regulates SEC16A levels and localization, specifically in TNBC. Altogether, we identified SEC16A as a key player in FER-driven TNBC invasion, highlighting the membrane trafficking machinery as a promising target for the development of new therapeutic strategies.

## Introduction

Triple-Negative Breast Cancer (TNBC) is the most aggressive subtype of breast cancer.^1^ Due to the absence of specific molecular targets, effective targeted therapies remain unavailable, and most patients ultimately succumb to metastatic disease. This underscores the urgent need of identifying and developing novel and effective therapeutics.

During the multi-step process of cancer cell invasion to surrounding tissue and metastasis to distant organs, adhesions between tumour cells and the Extra-Cellular Matrix (ECM) are altered to enable cell migration. This motile behaviour requires the endosomal recycling pathway, which controls the spatial distribution and turnover of adhesion components and signalling complexes.^2^ Endosomal sorting and recycling (ESR) is tightly regulated by constant Rab5-dependent endocytosis, followed by endosomal sorting that determines if proteins are either degraded or recycled back to the plasma membrane.^3^ Recycling activities involve Rab4, a GTPase that resides on recycling tubules emerging from early endosomes (EEs).^3^ Even though Rab5/Rab4 trafficking circuitry has been shown to be necessary for the spatial localisation of growth factor signalling driving invasive behaviour^4^, little is known about the differential regulation of endosomal sorting/recycling (ESR) in highly invasive versus poorly invasive cancer cells.

High expression of FPS/FES-related tyrosine kinase (FER) has been linked to tumour progression in several cancer types.^4–6^ In breast cancer, high FER expression is an independent predictor of decreased patient survival, especially in lymph node negative TNBC.^7^ Elevated FER levels correlate with high grade breast cancer types such as TNBC and breast cancer brain metastasis.^7,8^ Recently, we proposed that the FER expression levels could be used to predict patients’ response to taxane-based adjuvant chemotherapy.^9^ Importantly, our previous studies demonstrated that elevated expression of FER controls migration and metastasis formation by regulating the trafficking of recycling endosomal vesicles. Specifically, depletion of FER disrupts the formation of Rab4-positive recycling vesicles.^9^ Through modulation of the ESR pathway, FER influences the spatial distribution of α6/β1-integrins, thereby enhancing cellular invasion. Additionally, FER has been implicated in maintaining the localisation of growth factor receptors at the plasma membrane by orchestrating endocytosis and recycling processes.^10,11^ Thus, targeting the ESR pathway promoting the invasion of TNBC expressing FER could represent a promising targeted therapy. Yet, the FER-dependent ESR machinery involved in basal-like/TNBC invasion remains unclear.

Most of our knowledge on TNBC originates from studies using established cell lines in two-dimension (2D) settings. Even though these models have been invaluable to understand some aspect of TNBC pathogenesis, they are poorly suitable to study patient-specific tumour biology and harbour genetic alterations that may not be present in the original tumour. In contrast, three-dimension (3D) cultures of TNBC patient-derived xenograft organoids (PDXOs) recapitulate histological and genetic features of original tumours, including their intra-tumoral heterogeneity.^12,13^ In addition, PDXOs maintain the complex cell-ECM interactions, required for cell migration and invasion in 3D, which involves remodelling of tissue structures, exertion of pulling forces, and movement through Collagen I-rich matrix.^14^ Yet, little is currently known on the machinery promoting the invasion of patient-specific TNBC in 3D.

Here, we combined single-cell RNA sequencing, (phospho)proteomics and functional cell biology to identify the machinery driving the invasion in FER-expressing TNBC PDXOs. We identified membrane trafficking-regulators driving aggressiveness in TNBC, which may represent an opportunity for future targeted therapeutical intervention in FER-expressing TNBC.

## MATERIALS AND METHODS

### PDXOs culture

The TNBC PDXO #1 and #2 (1915 and 1963, respectively) were established and characterised by the Park Lab (McGill University, Canada)^15^.The PDXO line #1 (GCRC1915Tc) was derived from an Invasive Ductal Carcinoma-Not Otherwise Specified (IDC-NOS) female breast cancer patient with a basal molecular subtype. The patient had distant metastases and was treated with adriamycin/doxorubicin, cyclophosphamide, carboplatin and taxol/paclitaxel. The PDXO line #2 (GCRC1963T) is derived from an IDC-NOS female breast cancer patient with basal molecular subtype. The patient presented no axillary lymph node nor distant metastases and had not been treated.

PDXOs were grown in Complete or Reduced media. Reduced medium was prepared from Advanced DMEM/F12 reduced (12634028, Gibco), supplemented with 1% penicillin-streptomycin (pen/strep, 15070-063, Gibco), 1% GlutaMAX™ supplement (35050-061, Gibco), and 1% HEPES (15630080, Lonza). Complete medium consists of Reduced medium supplemented with 1.25 mM N-Acetyl-L-Cysteine (NAC; A9165-5G, Sigma), 1x B27 (17504001, Gibco), 50 µg/mLPrimocin (ant-pm1, Invivogen), 100 ng/mL Noggin (120-10C-50UG, Peprotech), 250 ng/mL R-spondin-3 (120-44-100UG, Peprotech), 10 mM Nicotinamide (N0636-100G,Sigma), 5 ng/mL FGF-7 (100-19-10UG, Peprotech), 20 ng/mL FGF-10 (100-26-25UG, Peprotech), 500 nM A83-01 (SML0788-5MG, Sigma), 37.5 ng/mL Neuregulin (100-03-50UG, Peprotech), 500 nM SB202190 (S7067-5MG, Sigma) and 5 ng/mL hEGF (AF-100-15-1MG, Peprotech).

PDXOs were thawed, kept in culture in Matrigel (3562, Corning) and passaged at 1:3. PDXO cell were mechanically and chemically dissociated and collected into a tube with ice-cold Reduced medium. Cells were pelleted by centrifugation at 1500 rpm at 4 °C for 5 min. The pellet was resuspended in warm TrypLE Express (25300062, Gibco), incubated at 37°C for 3 min and resuspended to achieve single-cell suspension. Following addition of 10 mL of cold Reduced medium and centrifugation at 1500 rpm at 4°C for 5 min, cells were re-plated in Matrigel at 37°C. To prevent cells from attaching to the bottom of the plate bottom, 40 µL drops of Matrigel–cell suspension were inverted and incubated for 45 min at 37°C until polymerized. Complete medium was subsequently added until the gels were fully submerged. Culture medium was refreshed every 2-4 days after passaging.

### Cell lines and culture conditions

MDA-MB-231 cells (MM231) were provided by Cell Lines Service (Eppelheim, Germany). STR typing of all cell lines was verified by PCR. Subconfluent cells were treated with 3.9 nM doxycycline (DOX) (D9891-1G, Sigma-Aldrich) for at least 5 days. Cells were cultured in Dulbecco’s modified Eagle medium (DMEM; 11039-047, Invitrogen), supplemented with 1% penicillin-streptomycin (15070-063, Invitrogen) and 10% fetal bovine serum (FBS) (16050-122, Invitrogen). Cells were cultured at 37°C with 5% CO2.

For 3D assays, MM231 cells were added to Matrigel (356231, Corning) at a density of 1000 cells/50 μL Matrigel. Droplets of 50 μL were added to flat bottom optical plastic 24-well plates (Corning). Plates were incubated for 45 min. at 37°C to allow the Matrigel to solidify, after which 500 μL normal growth medium was added. Cells were cultured for 7 days at 37°C.

### Molecular Biology

Inducible SEC16A-specific shRNA constructs (SEC16A-iKD) were generated using two shRNA oligo nucleotides designed and cloned as described previously ^16^. In short, shRNA targeting sequences directed against human SEC16A were chosen from the BROAD RNAi Consortium database (http://www.broadinstitute.org/rnai/trc) with the following ID’s: #TRCN0000246017 for shSEC16A#1 (sh1) and #TRCN0000246019 for shSEC16A#2 (sh2). Sh1 and sh2 target the sequences: #1-CGTGCTGTGGAGACGAGAAAT and #2-GATTACTATGCAAGCTATTAC, respectively. The EZ-Tet-pLKO-Puro vector^16^ (Addgene #85966) was co-digested with NheI (NEB, R3131S) and EcoRI (NEB, R0101S) and ligated to phosphorylated oligos. Ligations were performed using T4 Ligase (NEB, M0202S) with100 ng vector DNA and an 8:1 insert:vector molar ratio. Ligation reactions were transformed into Stbl3 (Life Technologies) chemically competent E. coli by heat-shock and bacteria plated on LB-agar plates containing 100 μg/mL ampicillin and incubated at 37 °C.

Full length SEC16AWT, SEC16A S1327A and SEC16A S1327D cDNAs resistant to SEC16A shRNA#1 were generated by introducing three silent mutations in the EGFP-SEC16A plasmid^17^ (pEGFP-SEC16A was a gift from David Stephens (Addgene plasmid #36155; http://n2t.net/addgene:36155; RRID: Addgene_36155)) via site-directed mutagenesis. To introduce the silent mutations and the S1327A and S1327D mutations, the following pairs of primers were used: SEC16A shResistant FW(5’-TGCCTGCCGCGTCCACGTGTTGTGGTGACGAAAAATGGGGAGATTGGAGGCCGCACCTC-3’) and SEC16A shResistant RV (5’-GAGGTGCGGCCTCCAATCTCCCCATTTTTCGTCACCACAACACGTGGACGCGGCAGGCA-3’); SEC16A S1327A FW(5’-CCTCGCTTCACGGGGGCTTTTGACGATGACCCCGATCCGCA-3’) and SEC16A S1327A RV (5’-TGCGGATCGGGGTCATCGTCAAAAGCCCCCGTGAAGCGAGG-3’); and SEC16A S1327D FW (5’-CCTCGCTTCACGGGGGATTTTGACGATGACCCCGATCCGCA-3’) and SEC16A S1327D RV (5’-TGCGGATCGGGGTCATCGTCAAAATCCCCCGTGAAGCGAGG-3’). Mutagenesis was performed using the PrimeSTAR Max DNA Polymerase Ver. 2 Premix (Takara, R047A). Each 50 µL PCR reaction contained 1× PrimeSTAR Max Premix, 0.4 µM forward primer, 0.4 µM reverse primer, 5% (v/v) DMSO, and 20 ng/µL plasmid DNA template. The final volume was adjusted with nuclease-free water. PCR amplification was carried out with a initial denaturation cycle at 95°C for 2 minutes, followed by 16 cycles at 95°C for 30 seconds, 56°C for 20 seconds, and 68°C for 30 seconds per kilobase of plasmid DNA (∼11.8 kb total). A final extension step was performed at 72°C for 5 minutes. To remove the methylated DNA template, 1.5 µL of DpnI (R0176S, New England Biolabs®) was added to each PCR reaction and incubated at 37°C for 16 hours. Following digestion, 1 µL of each DpnI-treated reaction mixture was transformed into Stbl3 (Life Technologies) chemically competent E. coli. Heat-shocked bacteria were plated on LB-agar plates containing 50 μg/mL kanamycin and incubated overnight at 37°C.

Diagnostic digestions were performed using AgeI (R0552S, New England Biolabs®) and/or NheI-HF (R3131S, New England Biolabs®) according to the manufacturer’s protocol. In addition, all clones were validated using Sanger sequencing to confirm the introduction of synonymous mutations conferring resistance to shRNA targeting SEC16A (Fig S2C-E) and the phosphorylation mutations (S1327A and S1327D) (Fig S2A and B).

### Transfection

MM231 cells were seeded into 6-well plates at a density of 2 × 10⁵ cells per well, approximately 24 hours prior to transfection, to achieve 70–80% confluency. Transfection was carried out using Lipofectamine 2000 (11668027, Invitrogen), following manufacturer’s instructions. Each well received 2 µg of plasmid DNA encoding GFP-tagged SEC16A variants: wild-type (WT), S1327A, or S1327D. Cells were incubated at 37 °C for 16 hours, after which the transfection medium was replaced with complete growth medium. At 72 hours post-transfection, cells were fixed and immunostained for subsequent microscopy analysis.

### Virus generation and cell transduction

Production and infection with lentivirus for inducible shRNA (iKD) constructs targeting SEC16A have been described previously.^18^ In short, lentiviral particles were produced in HEK293T cells using third-generation packaging constructs. Supernatant containing viral particles was harvested 48 hours after transfection and passed through a 45μm filter. MM231 cells were transduced overnight in the presence of 4 μg/mL polybrene (TR-1003-G, Sigma-Aldrich). DOX-inducible cell lines were treated for 4 days with 2 μg/mL Puromycin (Merck, 540411), refreshed every 2 days.

To transduce PDXO lines, PDXOs were dissociated into a single-cell suspension. The suspension was then aliquoted into 1.5 mL microcentrifuge tubes and spun down at 2000 rpm for 5 minutes. The medium was aspirated off and pelleted cells were resuspended in the lentiviral-polybrene mix, containing 500 µL of lentiviral-containing medium and 10 µg/mL polybrene. Spin infection was done at 600 g for 1 hour, and the plate was then incubated at 37°C. After a minimum of 6 hours, PDXOs were collected, spun down at 2000 rpm for 5 minutes, and plated in Matrigel at 37°C. To prevent cells from attaching to the plate bottom, 40 µL drops of Matrigel–cell suspension were inverted and incubated for 45 min at 37°C until polymerized. Complete medium was subsequently added until the gels were fully submerged. After two days, PDXOs were selected with 0.5 μg/mL puromycin for a week. After selection, organoids were treated with Doxycycline (Cat# D9891, Sigma-Aldrich) (2 µg/mL) for 3 days and knockdown efficiency was assessed by immunoblotting analysis.

### Collagen-I invasion assay

PDXOs were collected from Cultrex Basement Membrane Extract (BME; 3533-005-02, Trevigen) or Matrigel using 6 mg/mL Dispase (Gibco, 17105041) for 15 minutes, at 37°C. PDXOs were washed with cold Reduced medium. Collagen-I networks were produced using Collagen-I from rat tail (354236, Corning) at 2 mg/mL at pH 7.0-7.5. In short, Collagen-I was neutralised using 1N NaOH and further diluted using sterilised milliQ water and PBS 10x. Reduced medium containing PDXOs was added to the NaOH-neutralised solution (1/4 of the total volume). Hanging drops (40 µL) were made directly after neutralisation. Plates were inverted several times during the first 3-8 minutes to ensure homogeneous PDXOs distribution within the networks. Collagen-Igels were polymerised at 26°C for a minimum of 75 minutes before growth media (Complete or Reduced media) at 37°C were added. After this polymerisation step, plates were placed in the incubator at 37°C. The invading phenotype was assessed using Organoseg^19^ after 2 days of culture in Collagen-I.

### 3D morphology assessment

Brightfield images were acquired by using a 10× objective on a LEICA DMi1 Microsystem. Images were white balanced (after preliminary autoexposure). At least 5 images were acquired per chamber well, and at least two wells were imaged per condition. Three independent experiments were performed.

### Immunofluorescence Staining and Confocal Imaging

For immunofluorescence (IF) microscopy analysis, 2 × 10^4^ MM231 cells were seeded in No. 1.5 coverslips pre-coated with Poly-L-Lysine (P78920, Sigma-Aldrich). Cells were washed with 1X PBS (Biowest, L0615) and fixed in 4% paraformaldehyde (PFA, 28908, Thermo Scientific™) for 45 minutes at room temperature (RT). Following fixation, cells were washed twice with 1X PBS and permeabilised using 0.1% Triton-X-100 (T8787, Sigma-Aldrich) in 1X PBS for 10 minutes at room temperature (RT). Then, cells were quenched in 0.15% Glycine (pH 7.4) (A1067, ITW Reagents) for 10 minutes at RT, followed by blocking with 1% bovine serum albumin (BSA, A3294, Sigma-Aldrich) in 1X PBS for 10 minutes at RT. Primary antibodies were incubated overnight at 4°C in blocking solution. Coverslips were then washed three times with 1X PBS and incubated with Alexa Fluor 546/647-conjugated secondary antibodies (Jackson ImmunoResearch, 1:200) and Phalloidin-iFluor 488 (ab176753, Abcam, 1:1000) in blocking buffer for 1 hour at RT. After three washes with 1X PBS, cells were stained with DAPI (D3571, Invitrogen, 1:500) for 8 minutes at RT, washed again with 1X PBS and and mounted in Vectashield® Antifade Mounting Medium (H-1000, Vector Labs). Samples were then imaged using the Andor BC43 CF Benchtop spinning-disk confocal microscope (Oxford Instruments, UK), equipped with a 60x Plan Apochromat oil-immersion 1.42 NA objective. Fluorophores were excited with 405 nm (DAPI), 488 nm (GFP), 568 nm (TRITC), and 647 nm (Cy5) laser lines. Image capture was performed with a sCMOS camera (pixel size: 0.102 × 0.102 µm) controlled by Fusion software. Z-stacks were collected with a step size of 0.292 µm. Following acquisition, all image stacks were deconvolved with *Huygens Professional* version 25.04 (Scientific Volume Imaging, The Netherlands, http://svi.nl), using the *Standard* Deconvolution Express Profile to enhance spatial resolution and signal contrast.

For IF of PDXOs, PDXOs embedded in Collagen-I were fixed with 4% PFA (28908, Thermo Scientific™) for 30 minutes at RT (20°C) and washed 3 times with 1X PBS for 10-15 minutes each. Fixed samples were blocked using 10% normal goat serum (G9023-10ML, Sigma) in 0.3% Triton-X (Sigma-Aldrich) in 1X PBS for at least 1 hour. Primary antibodies were diluted in antibody buffer (0.3% Triton-X with 1% w/v BSA in 1X PBS) and incubated overnight at 4°C with shaking. The next day, samples were washed at least four times with 1X PBS for 10-15 min each and subsequently incubated with secondary Alexa Fluor 488/546/647-conjugated antibodies (Jackson ImmunoResearch, 1:200), together with DAPI (D9542, Sigma, 2 µg/mL), in antibody buffer at 4°C with shaking for at least 16 hours. For BrdU staining, PDXOs were incubated with 10 µm BrdU (10280879001, Roche) for 2 hours and fixed with 4% PFA. Then PDXOs were washed with 1X PBS before 2M HCl treatment for 80 min. After thorough washing of the PDXOs, we followed the previously described IF protocol. Images were acquired on a Leica SP5 confocal laser scanning microscope (Leica Microsystems) controlled by the Leica AF software (Leica Microsystems), using a 40× Plan Apochromat 1.10 NA water-immersion objective. Pinhole size was set to ∼1 Airy unit. Samples were illuminated by Diode 405 nm (DAPI), Argon 488 nm (Alexa Fluor 488), DPSS 561 nm (Alexa Fluor 546), and HeNe 633 nm (Alexa Fluor 647) laser lines. Emitted fluorescence was collected using photomultiplier tube (PMT) detectors. Z-stack images were captured at a pixel size of 0.22 × 0.22 µm with a step size of 7.51 µm. Sequential scanning was performed to avoid spectral bleed-through, with line and frame averaging applied to improve signal-to-noise ratio.

Antibodies used for IF of MM231 cells were: mouse anti-Rab4 (sc-271982, Santa Cruz, 1:50), mouse anti-Integrin-ß1 (610468, BD Biosciences, 1:50), mouse anti-Paxillin (PXN; 610051, BD Biosciences, 1:200) and rabbit anti-SEC16A (HPA0056 84, Protein Atlas Antibodies, 1:50). Antibodies used for IF studies of PDXOs were: mouse anti-FER (4268S, Cell Signaling, 1:200), rabbit anti-FER (NBP1-20089, Bio-Techne SRL, 1:200), rabbit anti-KRT14 (905301, BioLegend, 1:200), mouse anti-Rab4 (sc-271982, Santa Cruz, 1:200), rat anti-Integrin-ß1 (610468, BD Bioscience, 1:200) and mouse anti-BrdU (G3G4, Hybridoma Bank, 1:200).

### Electron Microscopy and immunogold labelling

For immunogold labelling, samples were fixed by immersion in 0.05% glutaraldehyde and 2% PFA in 0.1M sodium cacodylate buffer (pH 7.4) solution for 1 hour. The samples were then washed with 0.1 M sodium cacodylate buffer, and post-fixed for 1 hour in 1% osmium tetroxide in 0.1 M sodium cacodylate buffer. Afterwards, the samples were stained 30 minutes with aqueous 1% uranyl acetate, dehydrated, and embedded in Embed-812 resin. Ultra-thin sections (70 nm thick) were cut using an RMC Ultramicrotome with Diatome diamond knives, mounted on 200 mesh nickel grids. After washing the samples in TBS, they were incubated with Maxwell’s Solution for 5 minutes. This was followed by five washes with methanol and five washes with MilliQ water. The samples were then treated with 4% H₂O₂ for 5 minutes, followed by another five washes with MilliQ water. After heating at 95°C, the samples were washed with TBS containing 0.02% glycine for 10 minutes to quench free aldehyde groups.

Then samples were washed in TBS 0.5% BSA and incubated with TBS 1% BSA containing primary antibody: rabbit anti-Rab4 (ab13252, Abcam, 1:200) for 3 hours. The grids were then washed three times in TBS 0.5% BSA and incubated with 10 nm anti-rabbit IgG (whole molecule) gold conjugated (EM GAR10/1, BBI-Solutions OEM Limited, 1:20) for 1 hour at RT. The grids were then washed three times in TBS 0.5% BSA, incubated with 1% glutaraldehyde for 5 min and washed six times with deionised water prior to negative staining. Visualisation was carried out on a JEOL JEM 1400 TEM at 80 kV (Tokyo, Japan). Images were digitally recorded using a CCD digital camera PHURONA, EMSIS Germany at the Histology and Electronic Microscopy (HEMS) platform at i3S of the University of Porto.

### Single-cell isolation for mRNA-seq and flow cytometry analysis

PDXOs were digested using 2 mg/mL of Collagenase for a maximum of 5 minutes, at 37 °C. The single cell suspension was washed twice with pre-warmed Reduced medium. Cells were then resuspended in FACS buffer (0.1mM EDTA-20%FBS) and filtered using a 70-μm filter to remove debris. After centrifugation, the cells were resuspended in sorting buffer with 7-AAD (420403, BioLegend 1:300) to gate the viable cells. The sorting gating strategy is shown in Fig S2A. Fluorescence-activated cell sorting (FACS) was conducted using a FACS Aria III (BD Biosciences). The data were analysed using FlowJo (BD Biosciences, Fig S1A).

### Library preparation for scRNA-seq

The scRNA-seq libraries were generated from sorted viable single cells using Chromium Single Cell 3′ Reagent Kits v3.1 (10x Genomics, PN-1000127, PN-1000269, PN-1000215) according to the manufacturer’s instructions. The libraries were sequenced on a Novaseq 6000 sequencer (Illumina).

### scRNA-seq data analysis

Alignment of the scRNA-seq data was carried in Cell Ranger v7.1.0 using the GRCh38 human reference transcriptome. A count matrix table was generated for each sample, individually loaded into R (4.2.2) and analysed in Seurat v4.3.0. Only cells ranging from 2000-7000 unique features and less than 10% of expression of mitochondrial genes (to remove dead cells) were kept. Another common source of variation in scRNAseq data is the presence of doublets, therefore they were removed. After the quality control step, we were left with 20013 cells from PDXOs cultured in invasive condition (9983 invasive_1 and 10030 invasive_2) and 13661 cells from PDXOs grown in non-invasive condition (7148 non_invasive_1 and 6513 non_invasice_2) (Fig S1B).

After the quality control step, raw count data was normalised using the Seurat SCT transformation which performs normalisation, variance stabilisation, and feature selection based on a UMI-based gene expression matrix. Prior to this step, data belonging to the same condition were merged batch effects were found between cells from invasive and non-invasive conditions and normalized. To do so, we selected the 3000 most highly variable genes and removed unwanted sources of variation through regression of common contaminants such as the cell cycle stage and percentage of mitochondrial transcripts in the cell. Cell cycle scoring was calculated using the CellCycleScoring Seurat function, with the list of genes present in the S and G2M stages, given by Seurat. Integration was carried out by performing a canonical correlation analysis (CCA) to identify shared sources of variation between the datasets. Cell cycle gene analysis showed comparable number of cells in S-phase of the cell cycle between datasets (Fig S1C). Anchors (cells) were then identified and used to integrate the datasets. Variable genes between datasets were identified by the SelectIntegrationFeatures function, followed by PrepSCTIntegration and FindIntegrationAnchors to find the integration anchors. After this step, data were integrated using IntegrateData Seurat function.

Next, we performed the principal component analysis (PCA) and selected the first 30 PCs, based on both the inspection of PC elbow plot and through maxLikGlobalDimEst function from intrinsicDimension R package which estimates the intrinsic dimension of a data set using models of translated Poisson distributions).

Next, we applied the FindNeighbors function, that takes as input the previously defined dimensionality of the dataset (first 30 PCs) with k-nearest neighbour parameter of 20. To cluster the cells, the Leiden algorithm was used to interactively group cells together with the igraph method to avoid casting large data to a dense matrix, with the goal of optimising the standard modularity function. The clusters expressed comparable number of transcripts and expressed genes (Fig S1D and S1E). Invasive conditions enriched for cells belonging to the myoepithelial (446 cells in non-invasive condition versus 1123 cells in invasive condition) and basal cancer epithelial subpopulations (9002 cells in non-invasive condition versus 15057 cells in invasive condition), while the number of luminal cells were identical in both condition (1960 cells in non-invasive condition versus 1969 cells in invasive condition).

Then, we applied a cluster resolution of 0.4 for further analysis. Clustering analysis was carried using the t-Distributed Stochastic Neighbour Embedding (t-SNE) and the Uniform Manifold Approximation and Projection (UMAP) methods using the uwot method. Annotation of single-cell data was performed by manual identification of cluster. To do so, we followed the classifications made in^20^. The clusters were composed of different amounts of cells (Fig S1F), yet they expressed comparable number of transcripts and expressed genes (Fig S1G). Genes differentially expressed between clusters were detected using the FindAllMarkers function. A gene was only considered as differentially expressed if detected in a minimum percentage of 25% in either of the two groups of cells and having a minimum log2Fold change of 1. Only markers with an adjusted p-value below 0.05 were kept.

Over-representation analysis of the upregulated and downregulated genes in each cluster was carried using EnrichR with the GO_Molecular_Function_2018, GO_Biological_Process_2018, GO_Cellular_Component_2018, Azimuth_Cell_Types_2021, CellMarker_Augmented_2021, Descartes_Cell_Types_and_Tissue_2021. Only pathways with a p-value below 0.05 were recorded (Table S1 and S2).

Differential expression analysis between Invasive and non-invasive conditions was carried using the FindMarkers Seurat function for each identified cluster. Again, genes were only considered as differentially expressed if detected in a minimum percentage of 25% from cells cultured in either invasive or non-invasive conditions and having a minimum log2Fold change of 1. Only markers with an adjusted p-value below 0.05 were kept (Table S2).

The single-cell RNA sequencing data generated in this study have been deposited in the European Nucleotide Archive (ENA) under accession number PRJEB105431.

### Proteomics and Phospho-proteomics

24×10^4^ PDXOs (4.6 ×10^6^ cells) in Collagen-I were lysed with a buffer containing 8M Urea, 1M ammonium bicarbonate (ABC)-buffer, phosphatase inhibitor cocktail 2 and 3 (Roche), protease inhibitor cocktail (Roche), 10 mM tris(2-carboxyethyl)-phosphine (TCEP), 40 mM 2-chloroacetamide (CAA). Cell lysis was performed with a probe sonicator for 2 minutes at 80% amplitude and 0.5 second cycle time followed by 4-fold dilution with 5 mM CaCl2 and overnight digestion with trypsin (1:25). After digestion, the samples were concentrated and cleaned up using Sep-Pak C18 columns. The samples were enriched for phospho-peptides, after elution from Sep-Pak column using 1 ml Buffer B (60% Acetonitrile, 2% Trifluoroacetic Acid, 50mM Citric Acid), by addition of pre-washed TiO2 beads (10:1 beads:protein; ZirChrom Sachtopore-NP, 5um, 300A) and incubation for 5 minutes at 37 °C on a shaker. The supernatant was then incubated again on a second batch of TiO2 beads. After incubation, phospho-peptides bound to the TiO2 beads were centrifuged, and the supernatant was removed. The beads were then washed 3 times with wash buffer 1 (60% Acetonitrile, 0.5% TFA) using centrifugation to remove remaining non-phosphopeptides and once with wash buffer 2 (60% Acetonitrile, 0.1% TFA). After removing the supernatant, peptides were eluted with 500 µL 5% NH4OH for 10 minutes on a shaker at RT, separated from the beads by centrifugation and acidified with FA to 10% end concentration. Samples were concentrated on a Stagetip and after elution resuspended with 14 µL Buffer (water, 0.1% formic acid), half of which was used for Mass–Spectrometry (MS) analysis. The peptides were separated with a 240-minute gradient on an in-house made C18 capillary column (75 micrometer ID) using an easy nlc and directly sprayed into an Orbitrap Eclipse mass spectrometer (Thermo scientific), full scan resolution of 240k, a mass range of 400-1200 and two faims CVs (−45V,−65V) both with a cycle time of 1 second. HCD MS2 fragmentation was triggered at a 10 e4 threshold with a 0.7 Da isolation width window of the quadrupole and a 30% collision energy level. Readout in the Ion Trap at rapid mode had an AGC target of 10000. Five independent experiments were performed. The MS proteomics data have been deposited to the ProteomeXchange Consortium via the PRIDE partner repository^21^ with the dataset identifier PXD068815.

Raw files, both for proteome and phospho-proteome were analysed with MaxQuant [version 1.6.3.4] using the integrated Andromeda search engine. Carbamidomethyl (C) was set as the only fixed modification, variable modifications were set as Oxidation (M) and Acetyl (Protein N-term), with phosphorylation on serine, threonine, and tyrosine [Phospho (STY)] for the phospho-proteome dataset. Protein identification was performed against the UniProt Homo sapiens reference proteome (taxonomy ID: 9606) fasta file.

ProteinGroups and Phospho(STY)Sites output files were further processed in R [version 4.3.3] and Perseus [version 1.6.15.0]. Contaminants, reverse hits, and proteins identified in fewer than three out of four replicates in at least one condition were excluded and intensities were log₂-transformed. To assess for missing values in the differential enrichment analysis, missing values were imputed using a left-shifted gaussian distribution to account for missingness not at random (MNAR); for the linear regression, missing values were imputed with the minimum value for clearer visualisation.

Principal Component Analysis (PCA) was performed to evaluate the consistency among replicates. This was followed by scatter plots of the median replicate intensities to assess protein abundance. A linear regression model was fit into the data and proteins in the upper and lower 2.5% of residuals were selected as the most condition-specific outliers.

Differential enrichment analyses were computed using the DEP R package and significantly changing proteins were hierarchically clustered and explored as heatmaps. Gene Ontology enrichment analysis of the enriched proteins was performed using the clusterProfiler package in R for both full proteome and phospho-proteome datasets. To identify interactions between proteins and functional associations, protein–protein interaction networks of the enriched proteins were analysed in STRING database [version 12.0].

Proteins uniquely phosphorylated in invasive PDXOs to gene ontology terms related to ESR and membrane trafficking: Negative regulation of endocytic recycling (GO:2001136); Regulation of endocytic recycling (GO:2001135); Vesicle-mediated transport (GO:0016192); Endocytic recycling (GO:0032456); Positive regulation of endocytic recycling (GO:2001137)), Cellular component organization or biogenesis at cellular level (GO:0071840); Cellular component organization (GO:0016043); Cellular component biogenesis (GO:0044085); Organelle organization (GO:0006996)).

### Immunoblotting analysis

PDXOs embedded in Matrigel gels were collected after 7 days of culture. After adding 500 μL of cold 1X PBS (18912-014, Gibco), samples were centrifuged at 1200 rpm for 4 minutes to pellet PDXOs, and the supernatant was discarded. 30 μL of Laemmli Sample Buffer 2x (161-0737, Bio-rad) in β-Mercaptoethanol (M6250-100ML, Sigma-Aldrich) were added to each microcentrifuge tube and boiled for 10 minutes at 95 °C.

PDXOs embedded in Collagen-I gels were collected after two days of invasion assay and placed in microcentrifuge tubes on ice. After washing, supernatant was discarded. 30 μL of Laemmli Sample Buffer 2x (161-0737, Bio-rad) in β-Mercaptoethanol (M6250-100ML, Sigma-Aldrich) were added to each microcentrifuge tube, and samples were sonicated in BioruptorⓇ Plus (Diagenode) with high power mode for 6 cycles at 4 °C. Each cycle consisted of 30 seconds on and 30 seconds off. For MM231 lysates, cells were scraped in TRIS lysis solution containing inhibitors of protease (cOmplete Tablets, 4693159001, Roche) and phosphatase (PhosSTOP Tablets, 4906837001, Roche) and lysed for 20 minutes on ice. The cell lysates were then mixed with Sample Buffer, boiled for 5 minutes, and then centrifuged at 16,168 g for 30 minutes at 4 °C to clarify the mixture. After boiling at 65 °C for 10 minutes and at 95 °C for another 10 minutes, samples were stored at −20 °C or directly loaded on SDS-PAGE gels. For frozen samples, after thawing, the boiling step was repeated, and the lysates were centrifuged at 8000 rpm for 5 minutes.

Equal amounts of protein extracts in addition to the Precision Plus Kaleidoscope Standard molecular marker (1610395, Bio-Rad) were analysed by SDS-PAGE electrophoresis. After transfer, membranes were blocked with Bovine Serum Albumin (BSA) 5% (A7906, Sigma-Aldrich) in TBS-T (TBS with 0.1% Tween-20 (30P5927, Sigma-Aldrich) for 1 hour at RT. Membranes were then incubated overnight at 4 °C with primary antibodies: rabbit anti-SEC16A (A300-648A, Bethyl Laboratories, 1:500) or mouse anti-HSC70 (SC-7298,Santa Cruz, 1:8000) in 1% BSA in TBS-T. After washing with TBS-T, membranes were incubated with the IRDye® 680RD Donkey anti-Mouse (925-68072, LI-COR Biosciences) and the IRDye® 800CW Donkey anti-Rabbit (925-32213, LI-COR Biosciences) secondary antibodies. Signals were detected using the Odyssey® CLx Infrared Imaging System (LI-COR Biosciences).

### Cell cycle profiling

MM231 SEC16A iKD #1 and #2 cells treated with DOX for 3 days were fixed in 70% ethanol. Then, cells were washed and resuspended in 300 µL of 1X PBS, supplemented with 100 µg/mL RNAse A (19101, Qiagen) and 20 µg/mL Propidium Iodide (P4170, Sigma). Samples were incubated for 30 minutes in a 37°C water bath in the dark. Flow Cytometry was performed at a low flow rate using a BD Accuri C6 Flow Cytometer. The FlowJo, LLC software was used for quantification of the percentage of S-phase cells.

### *In vivo* chicken embryo chorioallantoic membrane (CAM) assay

On embryonic development day 9 (EDD9), 1 × 10^6^ TNBC PDXO #1 cells were inoculated per embryo into a 5 mm silicone ring under sterile conditions on top of the growing CAMs. At the endpoint (EDD16), CAM xenografted tumours were processed. Tumours’ morphology and invasion were characterised by haematoxylin and eosin (H&E) staining. According to the European Directive 2010/63/EU, ethical approval is not required for experiments using embryonic chicken, and the Portuguese law on animal welfare does not restrict the use of chicken eggs for experimental purposes.

### Multiplexed immunofuorescence

A five-plex multiplex immunofluorescence (mIF) panel was designed to characterize invasive TNBC PDXOs *in vivo*. The markers selected included FER, Sec16A, CD29, Rab4 and E-Cadherin. Formalin-fixed paraffin-embedded (FFPE) CAM tumour tissues (4 µm thickness) were used. All multiplex staining steps were fully automated and performed on the Ventana Discovery Ultra platform (Roche Diagnostics). Slides were subjected to automatic deparaffinization, followed by antigen retrieval using Cell Conditioning 1 (CC1) buffer (Roche Diagnostics, Cat# 950-124) for 40 minutes at 95 °C. Endogenous peroxidase activity was blocked using DAKO Hydrogen Peroxide Blocking Reagent (Agilent, Cat# S2023) for 20 minutes. Non-specific binding was blocked with Akoya Biosciences Blocking/Diluent Reagent (Cat# ARD1001EA) for 20 minutes. Primary antibodies were sequentially applied and incubated for 1 hour at 37 °C. The antibodies used were: anti-FER (4268S, Cell Signaling Technology, 1:2500), anti-SEC16A (HPA005684, Protein Atlas Antibodies, 1:100), anti-Integin-β1 (MA5-27900, ThermoFischer Scientific, 1:100), anti-Rab4A (MA5-17161, ThermoFischer Scientific, 1:1000), anti-E-cadherin (3195, Cell Signaling Technology, 1:50). Secondary detection was performed using the ultraview universal HRP multimer (Roche Diagnostics, Cat# 760-500) for 16 minutes. Signal amplification was achieved using Opal TSA fluorophores (Akoya Biosciences) diluted 1:300 in Opal Amplification Buffer (Cat# FP1498) and incubated for 32 minutes.. Between each staining round, antibody stripping was performed using CC2 (Roche Diagnostics, Cat# 950-223) buffer for 20 minutes at 100°C. The staining sequence and fluorophore assignment were as follows: FER (Opal 570), Sec16A (Opal 520), CD29 (Opal 480), Rab4 (Opal 620) and E-cadherin (Opal 690). Slides were counterstained with DAPI (Akoya Biosciences, Cat# FP1490) for 20 minutes and coverslipped using a fluorescence-compatible antifade mounting medium. Multispectral images were acquired using the PhenoImager HT system (Akoya Biosciences), running software version HT2.0 with integrated spectral unmixing capabilities. Whole-slide scans were acquired in MOTiF mode and exported in 8-bit QPTIFF format.

### Clinical samples and Immunohistochemistry (IHC)

The clinical sample sets and tissue microarray assembly have been previously described.^22,23^ Immunohistochemical staining was performed using the OptiView detection kit (Ventana, Roche diagnostics) on the Ventana Benchmark Ultra platform (Roche diagnostics). Tissue sections were baked at 75 °C for 8 minutes, then deparaffinized and heated back to 75 °C, followed by antigen retrieval using ULTRA Cell Conditioner #1 at 100 °C for 24 minutes. Anti-SEC16A primary antibodies (HPA005684, Protein Atlas Antibodies, 1:100) were manually applied and incubated at 37 °C for 32 minutes. Counterstaining was performed with Hematoxylin II for 12 minutes. All reagents were dispensed according to manufacturer’s instructions.

### Scoring of immunohistochemistry

All scorings were done by two individual observers blinded to patient characteristics and the outcome of other stainings. FER staining was scored previously.^7^ SEC16A was scored based on the intensity of the cytosolic staining, scores of 0 and 1+ were scored as low, and 2+ and 3+ were scored as high intensity.

### Quantifications

#### 3D Morphology Quantification

To assess cell invasion, the OrganoSeg software^19^ was used to streamline morphometric profiling from DIC images of 3D culture, acquired 2 days after invasion assay. The parameters were optimised manually for each image until a suitable segmentation was achieved. Invasiveness was inferred using the ‘Solidity’ parameter (Invasiveness=1-Solidity). The Prism 9.0.0 software (GraphPad) was then used to conduct the statistical analysis. Statistical significance was calculated using One-way ANOVA for PDXOs #1 and Mann-Whitney test for PDXOs #2. The data presented represent at least three biological replicates and are shown as mean ± Standard Deviation (SD).

#### Immunoblotting Analysis

The Image Studio™software (LI-COR Biosciences) was used to quantify the signal intensity of each band. Fold changes in SEC16A protein levels were determined by first normalizing the signal intensity of each SEC16A band in MM231-SEC16A iKD cells (cultured with or without doxycycline) to the corresponding HSC70 loading control. The SEC16A levels in cells cultured in plain medium were then normalized to the ones in the Doxycycline-containing medium.

#### Confocal Microscopy Image Analysis

To quantify the number and 3D spatial distribution of SEC16A puncta relative to the nucleus, Rab4-positive vesicles, and Paxillin-labelled focal adhesions in both MM231 FER-iKD and SEC16A-iKD cells, images were analysed using the Imaris 10.2.0 software (Oxford Instruments) (Fig S2F).

To quantify SEC16A distance to the nucleus in images from MM231 FER-iKD cells treated with or without DOX, nuclei were segmented from the DAPI channel using the *Surface* tool. To improve segmentation accuracy, a Gaussian filter was applied to reduce non-specific noise. Segmentation was then proceeded using the *Absolute Intensity Thresholding* method, with manual selection of the intensity range of interest. We applied the *Volume* filter to remove false positives. The shortest distance from each SEC16A spot to the nearest nucleus was calculated using the *Shortest Distance to Surface* statistical function.

To quantify Rab4 and Paxillin puncta in images from MM231 SEC16A-iKD cells treated with or without DOX, we used the Imaris *Spots* tool. In the *Spots Creation Wizard*, the *Quality* filter was applied to exclude low-intensity false positives, and *Distance to Surface* excluded spots within the nuclear volume, ensuring only cytoplasmic structures were analysed. For Paxillin-stained FAs, an additional *Z-position* filter restricted detections to cells’ basal planes. Analyses were conducted in batch mode to maintain consistent segmentation and measurement parameters across images. The *Number of Spots* function quantified Rab4 and Paxillin puncta.

To quantify Rab4 and Paxillin puncta in DOX-treated SEC16A-iKD MM231 cells expressing GFP-SEC16A-WT, -S1327A, or -S1327D, a different segmentation strategy was employed to enable analysis of individual cells (Fig S2F). For this approach, images were segmented using the deep learning–based model Cellpose-SAM.^24^ Segmentation was performed on three-channel (GFP, DAPI, Paxillin or Rab4) Z-stack images, using a Python 3 notebook (see Supplementary Material for code and parameters). Cellpose-SAM segmented each XY plane of the Z-stack independently, generating 2D regions of interest (ROIs). These ROIs were then stitched across adjacent planes into 3D objects based on their spatial overlap. The 3D segmentation output was imported into Imaris and converted to a *Surface* object. Segmented objects with a size below 2,66 × 10^5^ voxels, as well as those intersecting the image XY borders, were excluded to avoid artefactual and partial detections. Segmentation results were then reviewed and manually corrected to address mislabelled or improperly merged cells. Rab4, SEC16A, and Paxillin puncta were quantified as previously described. Following segmentation, for each image, cell surface, nucleus, and Rab4-vesicles and FAs were imported into a single *Cells* object. This made it possible to relate any structure located within a cell surface to that cell, and to measure their spatial relationships on a cell-by-cell basis (see Supplementary Material for representative images and associated *Cells* objects 3D renderings). Cells were classified as transfected or non-transfected based on the ratio of cytoplasmic to nuclear SEC16A spots, with values ≥0.8 indicating successful transfection (Fig S2G and H).

### Statistical Analysis

IHC statistical analyses were performed using IBM SPSS Statistics version 31.0 (SPSS Inc., Chicago, IL, USA). For clinical samples, associations between categorical variables were examined using Pearson’s w2-test.

For all other experiments, statistical analyses are described in the corresponding figure legends. Visualisation and statistical analyses were performed using GraphPad Prism 10.2.4. Plots for quantifications represent three independent biological replicates and are shown as mean ± SD. p < 0.05 was considered statistically significant.

## RESULTS

### FER-positive TNBC PDXOs invade in Collagen-I and accumulate Rab4 in invading cells

To dissect the FER-dependent endosomal recycling machinery promoting the invasion of TNBC cells in patient-relevant models, we searched for TNBC PDXOs cultured in 3D basement membrane extract (Matrigel) expressing FER. We identified two PDXO lines (hereafter designated as PDXO #1 and PDXO #2) that expressed high FER levels by western blot (Fig. S3A). When cultured in Matrigel with medium supplemented with growth factors (Complete medium), PDXO #1 (Fig. 1A) and PDXO #2 (Fig. S3B) remained compact, indicating that they did not invade. However, in 3D Collagen-I with Complete medium, both PDXO lines invaded as single-cells and as multicellular strands with tip cells expressing the basal epithelial marker KRT14 (Fig 1A and Fig S3B), which has been shown to be associated with the emergence of invasive leader cells in Collagen-I.^25^ The invasive behaviours of PDXO #1 and #2 in Collagen-I depended on the presence of growth factors, as both lines lost their ability to invade when grown in medium lacking NAC, B27, Primocin, Noggin, R-spondin-3, Nicotinamide, FGF-7, FGF-10, A83-01, Neuregulin, SB202190 and hEGF (hereafter designated as Reduced medium) (Fig 1B and Fig S3C). BrdU staining indicated that the lack of growth factors also significantly reduced the proliferation of PDXOs grown in Collagen-I that occurred exclusively in cells located in the organoid core (Fig. S3D). Hereafter, the culture of Collagen-I-embedded PDXOs grown in Reduced medium will be designated “non-invasive condition”, while those grown in Complete medium will be referred to as “invasive condition”. In agreement with a role of FER in promoting the invasion of TNBC cells, FER accumulated in invasive PDXO cell populations cultured in invasive condition (Fig 1C and Fig S3E). Quantification by western blot showed that FER levels increased by almost 2-folds in PDXOs cultured in invasive condition, compared to those grown non-invasive condition (Fig. 1D).

**Figure 1:**
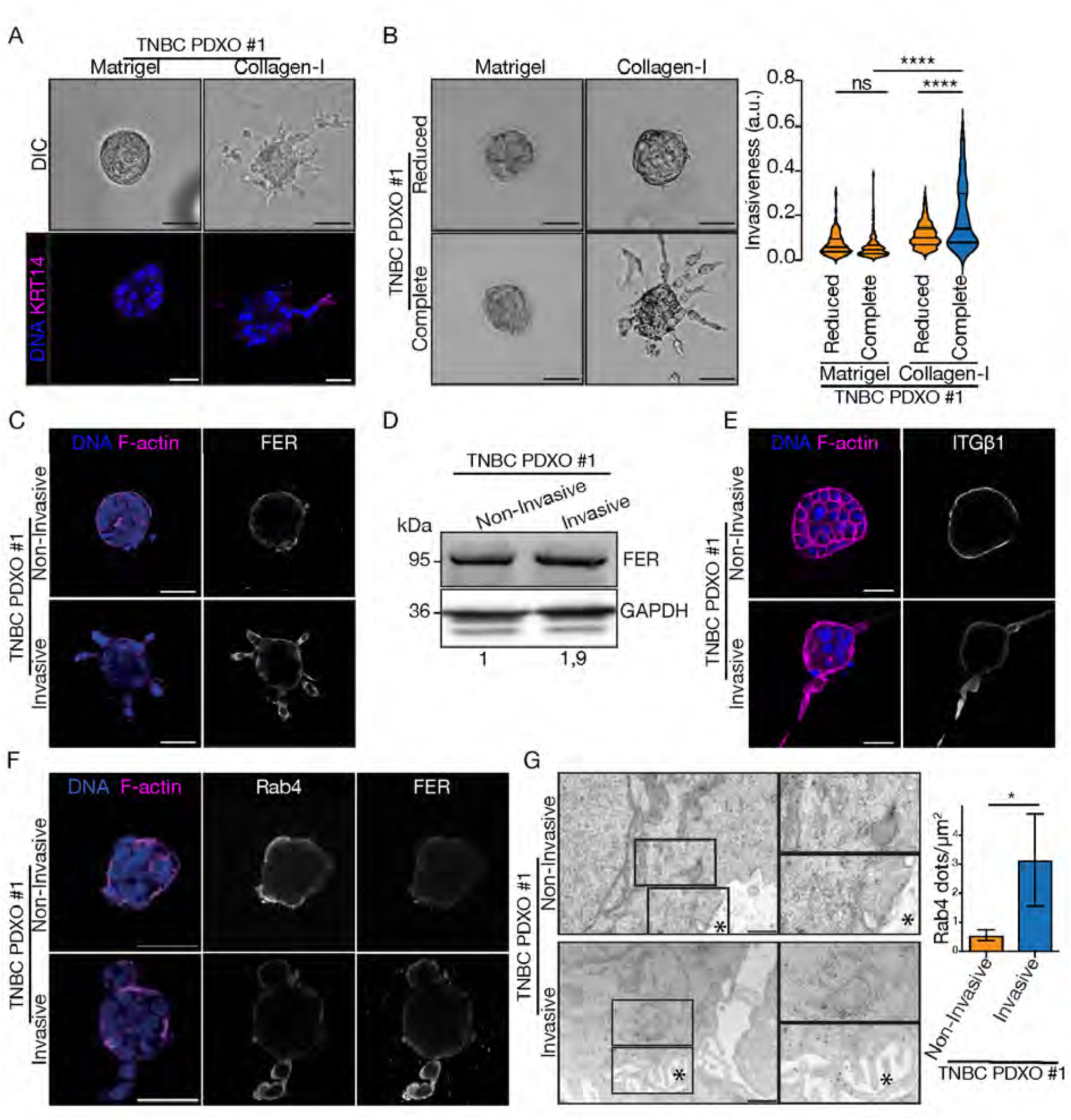
FER-expressing TNBC PDXOs #1 acquire invasive behaviours when cultured in Complete medium and Collagen-I. **(A)** DIC (upper panels) or confocal images (lower panels) of PDXOs #1 cultured in Complete medium and Collagen-I or Matrigel. (Lower panels) PDXOs are stained with DAPI (blue) to stain the nuclei and anti-KRT14 (magenta) to identify basal-like cells. Scale bars represent 50 μm. (**B)** (Left panels) DIC images of PDXOs #1 cultured in Reduced or Complete medium and Collagen-I or Matrigel. (Right panel) Quantification from three biological replicates of the invasiveness of PDXOs grown in the conditions indicated. More than134 PDXOs were quantified for each condition Statistical significance was calculated using one-way ANOVA. **** indicate p<0.0001. **(C)** Confocal images of PDXOs #1 cultured in Reduced or Complete medium and Collagen-I, stained with Phalloidin (magenta) to stain actin filaments (F-actin), DAPI (blue) and anti-FER (grey). Scale bars represent 50μm. **(D**) Western blot on protein extracts from PDXOs #1 cultured in Reduced or Complete medium and Collagen-I, blotted with anti-FER and anti-GAPDH used as a loading control. Numbers below indicate the fold-changes in FER levels for each condition. **(E)** Confocal images of PDXOs #1 cultured in Reduced or Complete medium and Collagen-I stained with Phalloidin (magenta), DAPI (blue) and anti-Integrin β1 (grey). Scale bars represent 50μm. **(F)** Confocal images of PDXOs #1 cultured in Reduced or Complete medium and Collagen-I, stained with Phalloidin (magenta), DAPI (blue), anti-Rab4 (grey, middle panels) and anti-FER (grey, right panels). Scale bars represent 50 μm. **(G)** Transmission Electron Microscopy images of PDXOs #1 cultured in Reduced or Complete medium and Collagen-I, stained for endogenous Rab4 (dense small black dots). Scale bars represent 0.5 µm. * indicates extracellular space. Inset images correspond to a 174% magnification. Quantification from three biological replicates of the number of Rab4 dots per μm2 for the conditions indicated. Error bars represent the standard deviation (SD). *p=0.0493. Statistical significance was calculated using unpaired t-test.

We reported that FER promotes the recycling of α6 and β1 integrins and the formation of Rab4 recycling tubules^7,9^. Consistent with these observations, β1 integrin was enriched in invasive PDXO cells grown in invasive conditions (Fig 1E). These cells also showed high Rab4 levels, which co-localised with FER (Fig 1F and Fig S3F). To assess Rab4 at subcellular level, we performed immuno-electron microscopy (IEM) by immunogold labelling of ultrathin cryosections of PDXOs. In invasive conditions, cells in contact with Collagen-I exhibited Rab4-positive structures that had a dense content and were often found in clusters (Fig 1G). Moreover, Rab4-positive signals could be found at the cytoplasmic membrane particularly at the tip of filopodia in invasive PDXO cells (Fig 1G), suggesting a fully functional recycling back to the membrane. Quantifications of Rab4-positive dots in PDXO cells in contact with Collagen-I indicated that invasive conditions triggered the accumulation of Rab4-positive dots, compared to non-invasive conditions (Fig. 1G). Taken together, these observations indicate that FER-expressing TNBC PDXOs are relevant models to dissect the FER-dependent endosomal recycling machinery promoting the invasion of TNBC cells.

### The phospho-proteome of invasive PDXOs is enriched for organelle biogenesis and organisation

To identify the membrane trafficking machinery of invading TNBC PDXO cell, we compared the full proteome of invasive and non-invasive PDXOs. Mass spectrometry (MS) analysis identified a total of 589 peptides significantly altered between both conditions, with 244 proteins accumulated in invasive PDXOs, against 345 in non-invasive ones (Fig 2A and Table S1). However. FER levels were not significantly altered between both conditions. Pathway enrichment analyses of all peptides with significantly different levels between both conditions indicated that proteins involved in actin cytoskeleton organization were less abundant in invasive PDXOs (Fig 2B and Table S1). On the other hand, a known regulator of active β1 integrins recycling, RABGAP1, crucial for cell adhesion, migration, and invasion in breast cancer cells^26^, accumulated in invasive PDXOs (Table S1). Because invasion was not associated with major alterations in the levels of protein associated with membrane trafficking, we asked if post-translational modifications of the trafficking machinery could play a role by exploring the phospho-proteomic landscapes of invasive and non-invasive PDXOs. MS analysis identified a total of 1589 phospho-residues on 1191 proteins (Fig 2C). Among those, 754 phospho-residues were only found in invasive PDXOs, while 729 were recovered exclusively in non-invasive PDXOs (Table S2). In addition, 106 residues were increased (53 phospho-residues) or decreased (53 phospho-residues) in invasive PDXOs (Table S2). Validating our approach, we found phosphorylation events in EGFR and SRC differentially regulated in invasive FER-expressing TNBC PDXOs. These are well established effectores of FER.^27–29^ Gene Ontology (GO) and pathway enrichment analysis of proteins uniquely or differentially phosphorylated between invasive and non-invasive PDXOs showed a significant enrichment of proteins involved in “organelle organisation” and cellular organisation or biogenesis” (Fig 2D; Table S2). A small set of proteins were commonly decreased in the full proteome and the phospho-proteome of invasive PDXOs. Among these, we found proteins involved in cell adhesion and actin cytoskeleton regulation (Table S2), two processes affected during invasion.^30^ Surprisingly, FER did not appear in our phospho-proteome. We then searched for gene ontology terms related to ESR and membrane trafficking in the list of phospho-proteins associated with invasive and non-invasive PDXOs (Fig 2E). 64 proteins were associated with ESR, organelle organization and biogenesis (Table S2). In particular, invasive PDXOs were enriched in phospho-proteins involved in vesicle endocytosis (e.g. CSNK1D and TRAPPC4), membrane binding and targeting of GAG proteins and coat protein II (COPII) vesicle coating, while depleted in SH3- and Myosin II-binding proteins and in components implicated in vesicle budding from the membrane, including SEC16A (Protein transport protein sec16 homolog A), SEC23A, and SEC31A (Fig 2E). Together, these observations suggest that phosphorylation-mediated regulation of membrane trafficking is a key mechanism promoting the invasion of TNBC PDXOs in Collagen-I matrices.

**Figure 2:**
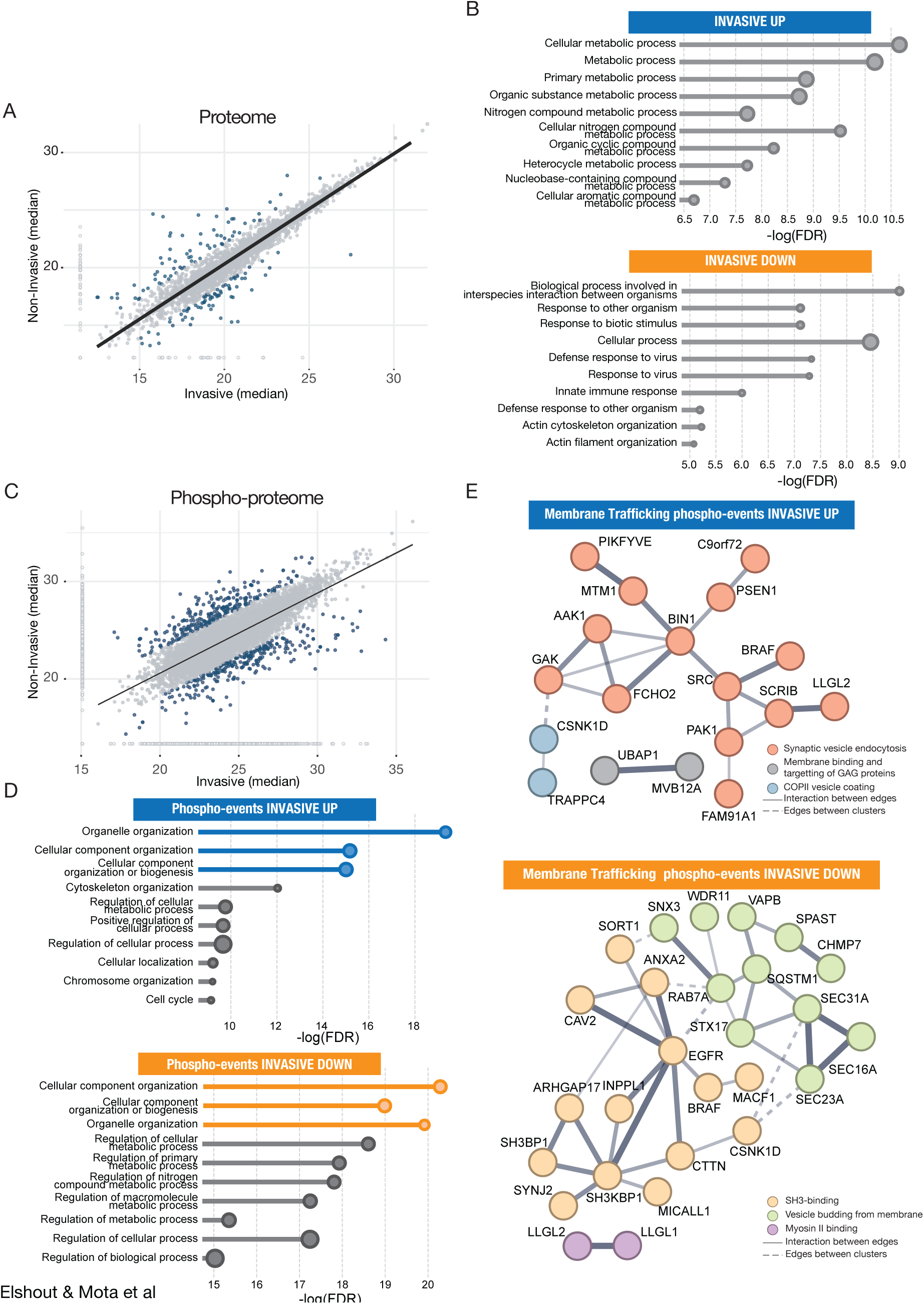
Invasive TNBC PDXO cells differentially phosphorylate regulators of membrane trafficking and organelle organization. **(A)** Scatter plot showing differentially regulated peptides between invasive (x axis) and non-invasive (y axis) PDXOs cultured in invasive and non-invasive conditions. **(B)** Pathway enrichment analysis of proteins presents at higher levels in invasive or non-invasive PDXOs. The 10 most affected pathways are plotted. Significantly altered pathways were filtered for adjusted FDR (<0.05). **(C)** Scatter plot showing differentially regulated phosphoresidues between invasive (x axis) and non-invasive (y axis) PDXOs cultured in Reduced or Complete medium and Collagen-I. **(D)** Pathway enrichment analysis of proteins phosphorylated in invasive or non-invasive PDXOs. The size of dots is proportional to the number of proteins identified. Significantly altered pathways were filtered for adjusted FDR (<0.05). Pathways of organelle organization and biogenesis are highlighted in colour. **(E)** Interaction network of proteins involved in ESR and organelle biogenesis, differentially phosphorylated in invasive and non-invasive PDXOs. Node colours represent functional clustering.

### The invasion of TNBC PDXO cells does not involve major transcriptional remodelling of membrane trafficking regulators

Although regulation of membrane trafficking in invasive TNBC PDXOs may not involve major alterations in proteins levels, our bulk proteomic analysis cannot exclude that the expression of membrane trafficking regulators is drastically affected in a minor population of invading cells. To assess the expression of genes associated with membrane trafficking in invading PDXO cells, we compared the expression profile of invading and non-invading FER-expressing TNBC PDXOs by single cell mRNA sequencing. Dimension reduction with UMAP unsupervised clustering analysis revealed three main cellular clusters (Fig. 3A and S4A). The major cluster (cluster 1) exhibited high levels of expression of the basal cell markers KRT5 and KRT14 (Fig 3A and B) and was defined as basal cancer cells. Cluster 2 expressed the epithelial tumour cells markers KRT18 and KRT19 (Cytokeratin 18 and 19), the proliferation marker MKI67 and the epithelial cells markers EPCAM (Epithelial Cell Adhesion Molecule) and lacked expression of KRT5 and KRT14. This cluster was therefore defined as a luminal subpopulation (Fig 3A and B). Cluster 3 displayed characteristics of a myoepithelial cell population, as it expressed high levels of KRT5 and KRT14 (Fig 3A and B).

**Figure 3:**
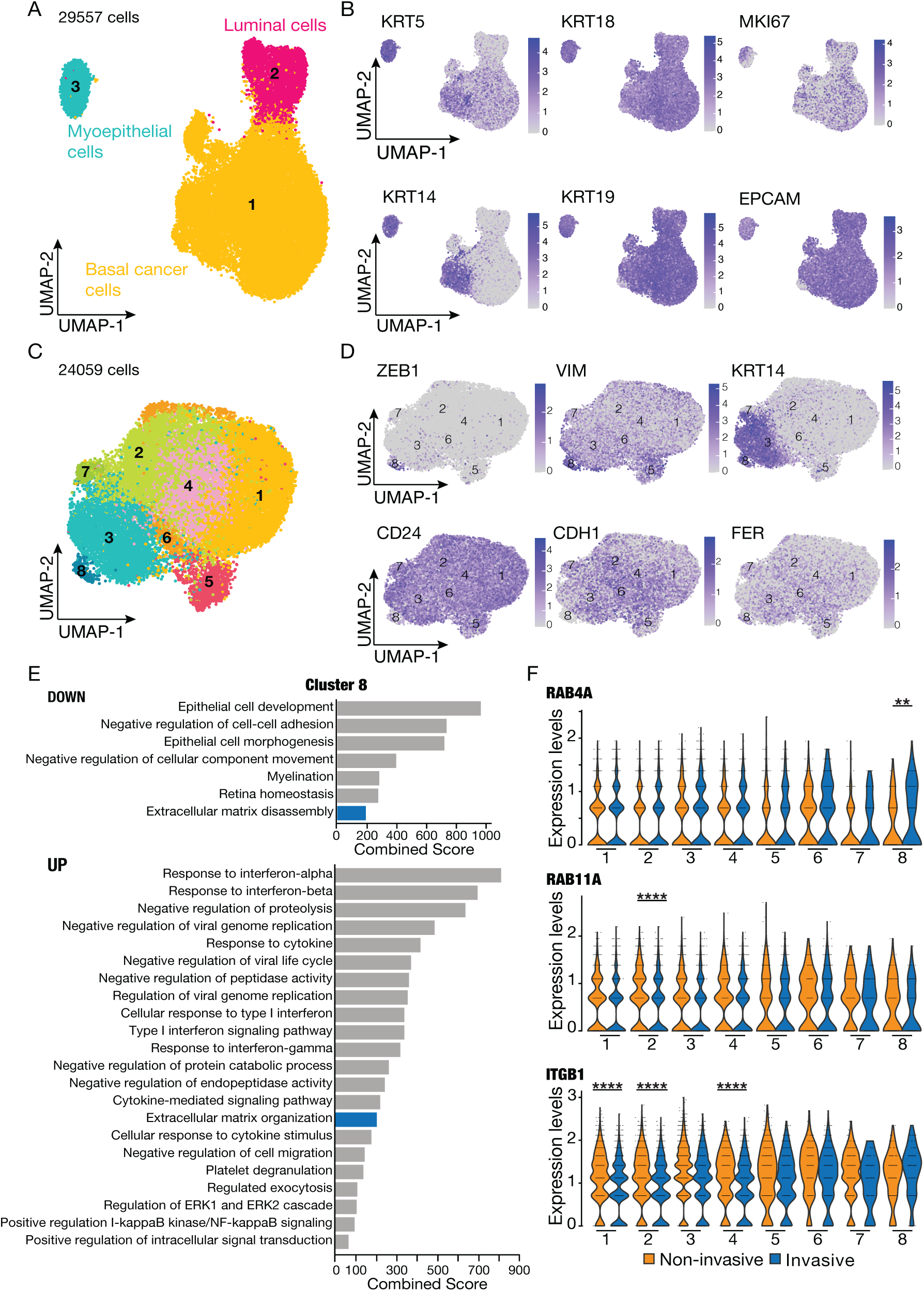
The membrane trafficking machinery is not primarily regulated at transcriptional levels in invasive TNBC PDXO cells. **(A)** UMAP visualisation of 29,557 TNBC PDXO cells cultured in invasive or non-invasive conditions and analysed by scRNA-seq. Clusters annotated for their cell types, predicted using canonical markers and signature-based annotation, identify a Myoepithelial cell population (cyan), a Luminal cell population (red) and a Basal cancer cell population (orange). **(B)** UMAP of PDXO #1 cells indicating the log-normalized expression of markers identifying basal-like cells (KRT5 and KRT14), epithelial tumour cells (KRT18 and KRT19), proliferative cells (MKI67) and epithelial cells (EPCAM). **(C)** UMAPs of the 8 clusters constituting the basal cancer cell population of PDXOs cultured in invasive or non-invasive conditions (24059 cells). **(D)** UMAPs of the basal cancer cell population of PDXOs indicating the log-normalised expression of ZEB1, VIM, KRT14, CD24 and CDH1 and FER. **(E)** Pathway analysis of genes differentially expressed between cluster 8 and cluster 1 to 7 in PDXOs cultured in invasive and non-invasive conditions. Cluster 8 differentially expressed ECM components (blue). Significantly altered pathways were filtered for adjusted p-value (<0.05). **(F)** Quantification of the expression levels of RAB4A, RAB11A and ITGB1 in all clusters between PDXOs cultured in invasive (blue) and non-invasive (yellow) conditions. The y axis represents the distribution of expression through a density plot. Differential expression analysis between cells cultured in invasive and non-invasive conditions was carried using the FindMarkers Seurat function for each identified cluster. \*\**P*= 0.01, \*\*\*\**P* < 0.0001.

Because invading TNBC cells should belong to the basal cancer epithelial cluster, we further refined cell populations within cluster 1 based on differential expression profiles. By applying a resolution of 0.4 we identified 8 clusters (Fig 3C and S4B). To identify the invasive cell cluster among those, we searched for mesenchymal markers differentially expressed between all clusters. ZEB1, VIM and KRT14 were upregulated in cluster 8. In addition, this cluster downregulated the expression of CD24 and CDH1, which identify differentiated and epithelial cells, respectively (Fig 3C and D). Cluster 8 also downregulated RAB11FIP1 and RAB25 (Table S2), two RAB proteins associated with slow-recycling, which has been shown hindered in pro-invasive cells.^9,31,32^ However, although FER was slightly increased in clusters 7 and 8, these differences were not significant (Fig 3D and S4C). Strikingly, gene enrichment analysis for each cluster indicated that cluster 8 was the only cluster in which regulators of the ECM were differentially expressed (Fig 3E and Table S2).^9,31,32^ Moreover, this cluster downregulated regulators controlling an epithelial state. Furthermore, as expected for an invading cell population, cluster 8 was the smallest in term of cell number (Total: 272 (1,13% of all basal cancer cells); Invasive – 146 (0,97% of invasive basal cancer cells) and non-invasive – 126(1.4% of non-invasive basal cancer cells) (Fig S1G and Fig S4B). Thus, cluster 8 could encompass the TNBC cell population actively remodelling the ECM at the outer of PDXOs with pro-invasive abilities.

We then compared genes differentially expressed in cluster 8 between invasive and non-invasive PDXOs. Genes involved in membrane trafficking or organelle organization were not differentially expressed in this cluster between both conditions. Most pathways differentially expressed were related to viral-related processes and oxidative stress (Fig S4D). Moreover, neither RAB11, a regulator of slow recycling, nor ITGB1, were differentially expressed in cluster 8 between both conditions (Fig 3F). However, RAB4A expression was significantly increased by 1.57-fold in cluster 8 from invasive PDXOs, compared to the non-invasive cluster 8 (Fig 3F and Table S3). Thus, the upregulation of RAB4A in these cells could be required for invasion. Yet, the control over membrane trafficking, which triggers the invasion of TNBC cells, may not be preferentially regulated at transcriptional levels.

### SEC16A is necessary for invasion in TNBC cells

Expression of the endoplasmic reticulum (ER) export factor SEC16A was not significantly affected in cluster 8 between invasive and non-invasive conditions (Fig. S4C). Yet, invasion could require differential phosphorylation of SEC16A. The phosphorylation of SEC16A at Serine 1327 (S1327) is triggered by FER depletion in MM231 cells^9^ and also turned up in non-invasive PDXOs. To investigate if SEC16A play a role in basal-like breast cancer invasion, we generated two MM231 cell lines stably infected with independent doxycycline (DOX)-inducible SEC16A short hairpin RNA (MM231-SEC16AiKD). MM231-SEC16AiKD #1 (Fig 4A) or MM231-SEC16AiKD #2 (Fig S5A) cells treated with DOX showed efficient depletion of SEC16A, compared to cells grown in the absence of DOX (-DOX) by western blots. Consistent with previous reports^33^, SEC16A localized at the perinuclear region in untreated MM231-SEC16AiKD cells. This signal was no longer detected in DOX-treated cells (Fig 4B and Fig S5B). To test the effect of SEC16A depletion on the invasive potential of MM231 cells, we grew MM231-SEC16AiKD spheroids in Matrigel. Untreated MM231-SEC16AiKD (Fig 4C and Fig S5C) or DOX-treated MM231(Fig S5D) spheroids acquired a stellate morphology, reflecting their invasive capacity in Matrigel. In contrast, MM231-SEC16AiKD spheroids treated with DOX remained significantly rounder (Fig 4C and S5C), indicating that SEC16A is required for the invasion of MM231 cells. In agreement with previous reports^34^, SEC16A also impacted on cell proliferation, as DOX treatment significantly decreased the percentage of MM231-SEC16AiKD cells in S-phase of the cell cycle, compared to those left without treatment (Fig S5E). We then tested if SEC16A was also required for the invasion of TNBC PDXO cells by generating PDXOs stably infected with DOX-inducible SEC16AiKD #1. DOX-treatment caused a 50% reduction in total SEC16A protein levels, compared to untreated PDXOs carrying SEC16AiKD #1 (Fig 4D). Importantly, it resulted in a significant reduction in the invasiveness of PDXOs cultured in invasive condition, compared to those left without treatment (Fig 4E). This effect was not caused by DOX *per se,* as DOX-treated parental PDXOs invaded as efficiently as untreated PDXOs (Fig S5F). We then investigated if the inability of SEC16A-depleted cells to invade was associated with defects in adhesion and endosomal trafficking. DOX-treated MM231-SEC16AiKD cells showed a significant decrease in the number Paxillin foci, which marks Focal Adhesions (FAs), compared to untreated MM231-SEC16AiKD cells (Fig 4F), indicating that SEC16A is required for the assembly or maintenance of FAs. In addition, SEC16A-depleted cells displayed a marked decrease in Rab4-positive tubules relative to those expressing SEC16A (Fig 4G), suggesting that SEC16A stimulates Rab4-mediated endosomal transport. Together, these observations indicate that SEC16A is required for the invasion of basal-like breast cancer cells, likely by promoting the recycling of FAs.

**Figure 4:**
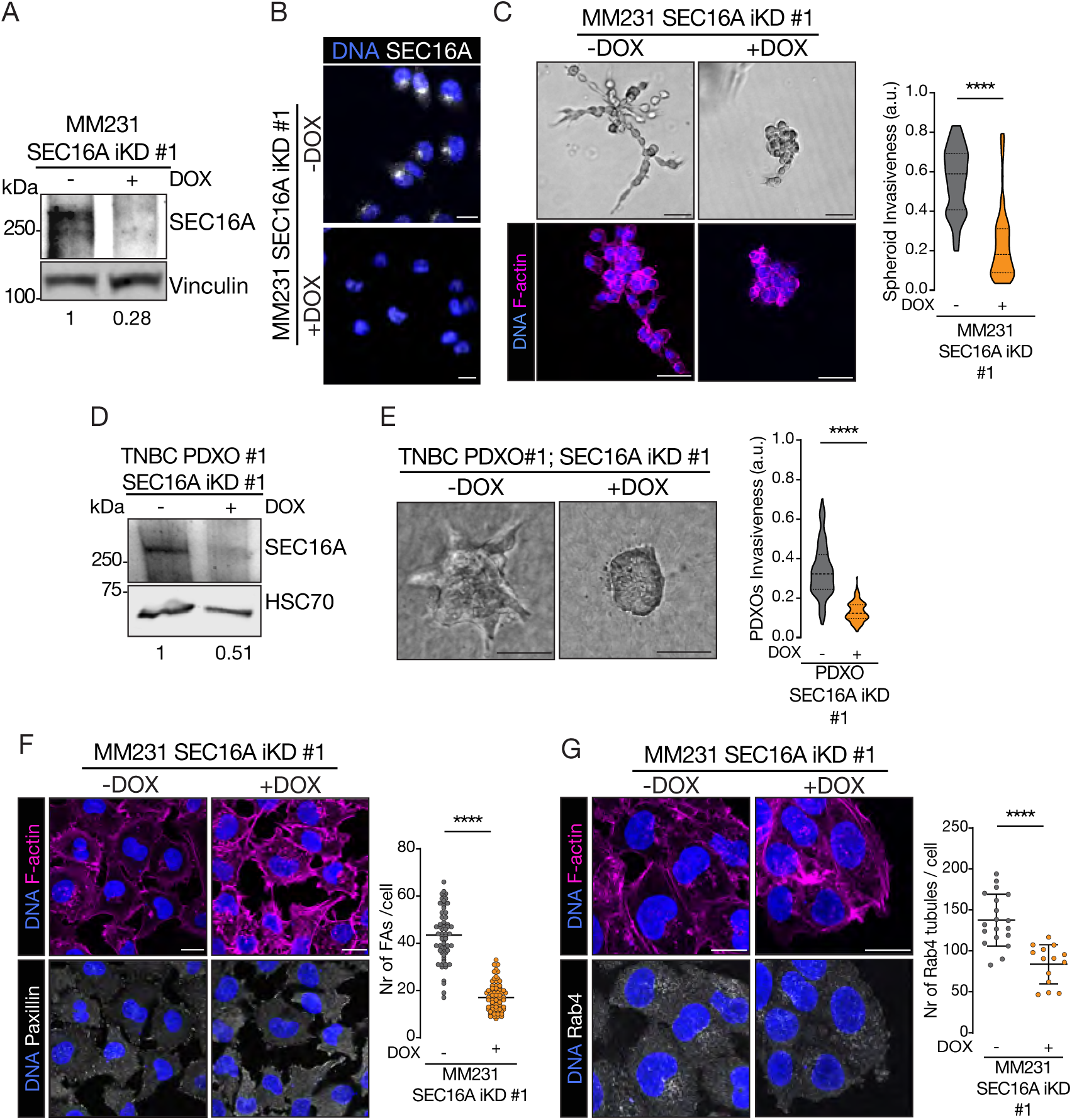
SEC16A depletion prevents TNBC invasion and reduces the number of FAs and Rab4 vesicles. **(A)** Western blot of protein extracts from MM231 SEC16A-iKD cells with or without DOX treatment, blotted with anti-SEC16A and anti-Vinculin used as loading control. Numbers on the bottom indicate the fold-changes in SEC16A levels normalised to loading control levels for each experimental condition **(B)** Confocal images of MM231 SEC16A-iKD cells untreated or treated with DOX, stained with DAPI (blue) and anti-SEC16A (grey). Scale bars indicate 15 µm. **(C)** (Left panels) Phase contrast (upper panels) or confocal images (lower panels) of MM231 SEC16A-iKD #1 spheroids in Matrigel untreated or treated with DOX. (Lower panels) cells were stained with DAPI (blue) and Phalloidin (magenta). Scale bars, 50 μm. (Right panel) Quantification of the invasiveness of MM231 SEC16A-iKD #1 spheroids untreated or treated with DOX. Statistical significance was calculated using unpaired t-test ****p<0.0001. **(D)** Western blot on protein extracts from TNBC PDXOs SEC16A-iKD untreated or treated with DOX, cultured in invasive conditions and blotted with anti-SEC16A and anti-HSC70 used as a loading control. Numbers on the bottom indicate the fold-changes in SEC16A levels normalised to HSC70 levels for each experimental condition. **(E)** (Left panels) DIC images of TNBC PDXOs SEC16A-iKD cultured in invasive conditions, untreated or treated without DOX. (Right panel) Quantification of the invasiveness of TNBC PDXOs SEC16A-iKD cultured in invasive condition, untreated or treated without DOX. Data are from three biological replicates and include more than 30 PDXOs per condition and per replicate. Statistical significance was calculated using unpaired t-test. ∗∗∗∗p<0.0001. **(F)** (Left panels) Confocal images of MM231 SEC16A-iKD cells untreated or treated with DOX, stained with DAPI (blue), Phalloidin (magenta), and Paxillin (grey). Scale bars indicate 15 µm. (Right panel) Quantification of the number of FAs per MM231 SEC16A-iKD cells treated with DOX (n=71) or without DOX (n=64). Data are from three biological replicates with at least three fields analysed per condition and per replicate. Each dot represents one cell. Statistical significance was calculated using unpaired t-test ****p<0.0001. **(G)** (Left panels) Confocal images of MM231 SEC16A-iKD cells untreated or treated without DOX, stained with DAPI (blue), Phalloidin (magenta) and anti-Rab4 (grey). Scale bars indicate 15 µm. (Right panel) Quantification of the number of Rab4 vesicles in MM231 SEC16A-iKD cells treated with DOX (n=336) or without DOX (n=496). Data are from three biological replicates, with at least three fields analysed per condition and per replicate. Each dot represents one field. Scale bars indicate 15 µm. Error bars indicate standard deviation. Statistical significance was calculated using unpaired t-test. ****p<0.0001.

### Phosphorylation of SEC16A could regulate endosomal recycling and FAs in TNBC cells

FER^7,9,35^ and SEC16A depletion have identical effects on the invasion of basal-like breast cancer cells. SEC16A is phosphorylated at S1327 in FER-depleted MM231^9^ and in non-invasive TNBC PDXOs (Fig 5A). Thus, SEC16A inactivation through S1327 phosphorylation could impair ESR-dependent cell invasion. To investigate the impact of SEC16A S1327 phosphorylation, we transfected short hairpin-resistant GFP-tagged versions of wild type SEC16A (GFP-SEC16A WT) or SEC16A mutant versions, either phospho-deficient (GFP-SEC16A S1327A) or phospho-mimetic (GFP-SEC16A S1327D) in MM231-SEC16AiKD cells. Alike endogenous SEC16A, GFP-SEC16A-WT, GFP-SEC16A-S1327A and GFP-SEC16A-S1327D localised at the perinuclear region in DOX-treated MM231 SEC16AiKD (Fig. 5B). Surprisingly, while the number of Rab4-positive structures were not significantly different between SEC16A-depleted cells expressing GFP-SEC16A-WT and those expressing GFP-SEC16A-S1327D, expressing GFP-SEC16A-S1327A in SEC16A-depleted cells reduced the number of Rab4-positive structures (Fig 5C). Similarly, SEC16A-depleted cells expressing GFP-SEC16A-S1327A had fewer Paxillin-positive foci, compared to those expressing GFP-SEC16A-WT or GFP-SEC16A-S1327D (Fig 5D). These observations suggest that S1327 phosphorylation is not required for the perinuclear localisation of SEC16A, but promotes the recycling of FAs. When analysing the protein region in which this residue sits (region 1307-1378), we observed that this sequence is enriched in phosphorylatable residues compared to the average human protein composition (23.61% vs. 15.9%, respectively)^36^, with 2 tyrosine (Y;2.63%), 1 threonine (T;1.32%) and 15 serine (S; 19.74%) (Fig 5E). It has been described that multiple phosphorylation events might have a direct impact in phosphorylation detection, as they can increase peptide’s negative charge and hydrophilicity, thereby reducing ionization efficiency in positive ion mode and impairing its detection by MS.^37^ So, if phospho-S1327 detection would be affected by the negative net charge of the peptide, it would only be enriched or detectable in conditions where the neighbouring residues are unphosphorylated. If so, S1327 phosphorylation could take place before other SEC16A phosphorylation events in the same region, and thus become undetectable under invasive conditions. Altogether, these findings suggest that phosphorylation of SEC16A at S1327 is required for endosomal recycling and FA assembly or maintenance, thereby enabling the invasive behaviour of FER-expressing TNBC cells.

**Figure 5:**
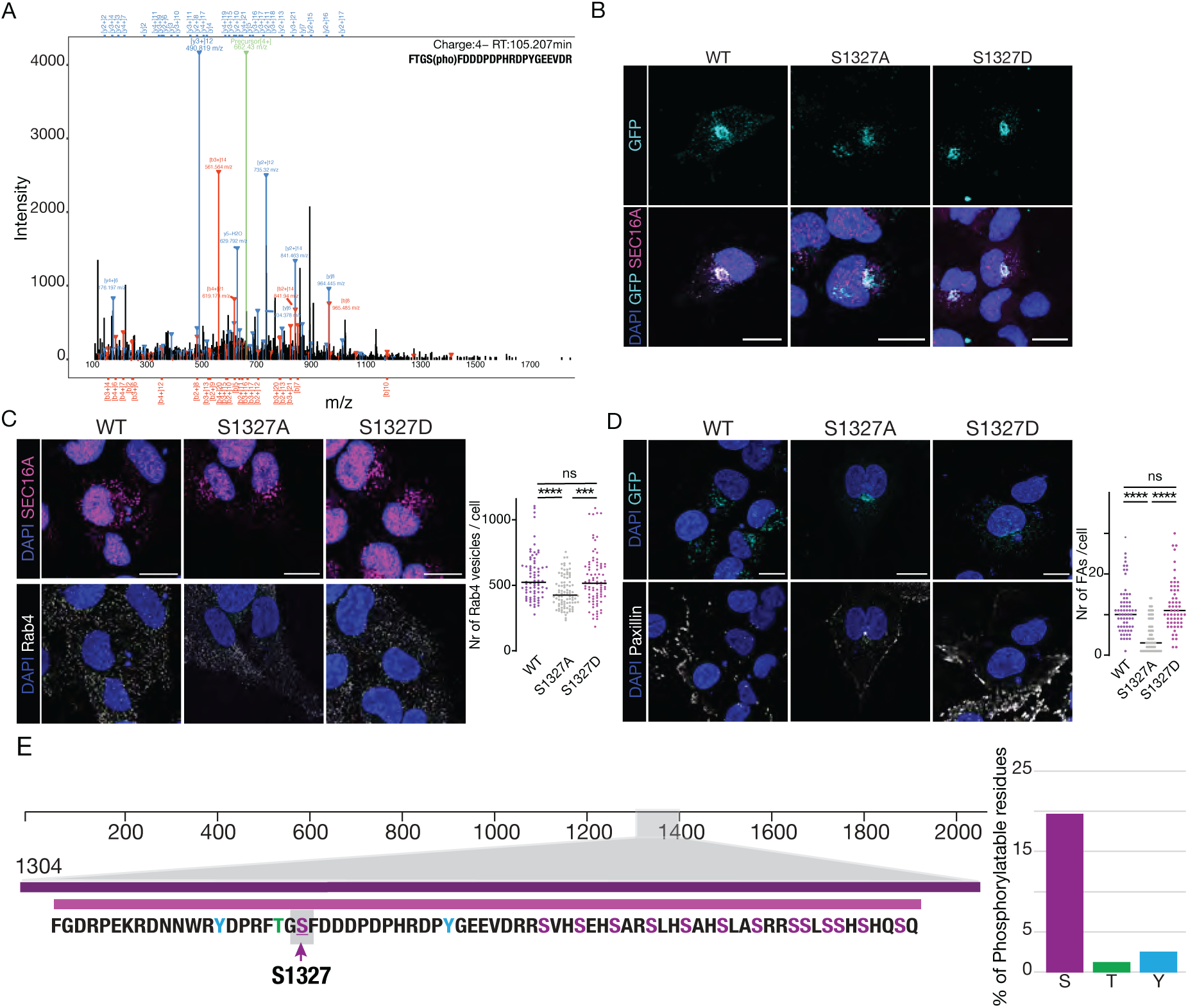
Phosphorylation of SEC16A regulates the number of Rab4 vesicles and FAs in TNBC cells. **(A)** MS/MS spectrum of the SEC16A phosphopeptide FTGS[pho]FDDDPDPHRDPYGEEVDR (4+), showing phosphorylation at Ser4, corresponding to Ser1327 in the full-length protein. Labelled peaks correspond to the main fragment ions observed in the spectrum. Identified b- and y-series ions, together with the precursor ion, confirm the peptide sequence. The b- ions identified support the presence and location of the phosphate group. The site was identified with a localization probability of 0.92 and a PEP score of 0.0085. **(B)** Confocal images of MM231 SEC16A iKD cells expressing GFP-SEC16A-WT (cyan blue), or GFP-SEC16A-S1327A (cyan blue) or GFP-SEC16A-S1327D (cyan blue), treated with DOX and stained with DAPI (blue) and anti-SEC16A (magenta). Scale bars represent 15μm. **(C)** (Left panels) Confocal images of MM231 SEC16A iKD cells expressing GFP-SEC16A-WT, or GFP-SEC16A-S1327A or GFP-SEC16A-S1327D, treated with DOX and stained with DAPI (blue), anti-SEC16A (magenta) and anti-Rab4 (grey). Scale bars represent 15μm. (Right panel) Quantification of the number of Rab4 vesicles in MM231 SEC16A iKD cells expressing GFP-SEC16A-WT (n=76) or GFP-SEC16A-S1327A (n=93) or GFP-SEC16A-S1327D (n=85). Data are from three biological replicates with at least three fields analysed per condition and per replicate. Each dot represents one cell. ****p<0.0001 between cells expressing WT SEC16A and S1327A SEC16A. ***p=0.0004 between cells expressing S1327A SEC16A and S1327D SEC16A. ns indicates non-significant between cells reconstituted with WT SEC16A and S1327D SEC16A (p=0.9291). Statistical significance was calculated using one-way ANOVA. **(D)** (Left panels) Confocal images of MM231 SEC16A iKD cells expressing GFP-SEC16A-WT (cyan blue), or GFP-SEC16A-S1327A (cyan blue) or GFP-SEC16A-S1327D (cyan blue), treated with DOX and stained with DAPI (blue) and anti-Paxillin (grey). Scale bars represent 15 μm. (Right panel) Quantification of the number of FAs in MM231 SEC16A iKD cells expressing GFP-SEC16A-WT (n=65) or GFP-SEC16A-S1327A (n=78) or GFP-SEC16A-S1327D (n=59). Data are from three biological replicates with at least three fields analysed per condition and per replicate. Each dot represents one cell. ****p<0.0001 between cells expressing WT SEC16A and S1327A SEC16A and between cells expressing S1327A SEC16A and S1327D SEC16A. ns indicates non-significant (p=0.4673) between cells expressing WT SEC16A and S1327D SEC16A. Statistical significance was calculated using one-way ANOVA. **(E)** (Left panel) Aminoacid sequence of the region comprising amino-acids 1307-1378. Phosphorylatable residues are highlighted: serine (magenta), threonine (green), or tyrosine (blue). Purple arrow indicates Ser1327. (Right panel) Ratios of serine (magenta), threonine (green), or tyrosine (blue) residues in the region comprising amino-acids 1307-1378.

### FER controls SEC16A subcellular localisation and levels in basal-like breast cancer cells

We next investigated whether FER regulates SEC16A. Due to the lack of SEC16A phospho-specific antibodies, we assessed the effect of FER on SEC16A levels and localisation in MM231 cells stably infected with DOX-inducible short hairpin RNA against FER (MM231-FERiKD). Surprisingly, while FER levels were reduced in DOX-treated MM231-FERiKD, SEC16A levels were almost increased by 2-folds (Fig 6A). In addition, FER controls SEC16A subcellular localisation, as the perinuclear accumulation of SEC16A in untreated MM231-FERiKD was lost when cells were treated with DOX (Fig 6B). These observations indicate that FER promotes SEC16A localisation at the perinuclear region, and suggest that the lack of SEC16A perinuclear localisation restricts SEC16A protein turnover. To address if FER also controls SEC16A levels and localisation *in vivo*, we xenografted TNBC PDXO #1 cells on chicken embryo chorioallantoic membrane (CAM). PDXO cells were able to give rise to tumour mass and exhibited invasive behaviour *in vivo*. Multiplex immunohistochemistry of PDXO tumours on CAM indicated that FER, Rab4, and Integrin β1-positive cells were enriched at the invasive margin (Fig. 6C). Consistent with the downregulation of E-cadherin (E-cad), encoded by CDH1, in cluster 8 (Fig. 3D), a close up of invasive PDXO cells indicated that E-cad levels were markedly reduced in invasive leader cells, compared to inner tumour cells (Fig. 6D). Also, leader cells accumulated FER, while shown lower levels of SEC16A. In contrast, follower cells that exhibited lower FER levels, accumulated SEC16A (Fig 6D). Thus, *in vitro* and *in vivo*, FER and SEC16A levels are inversely correlated.

**Figure 6:**
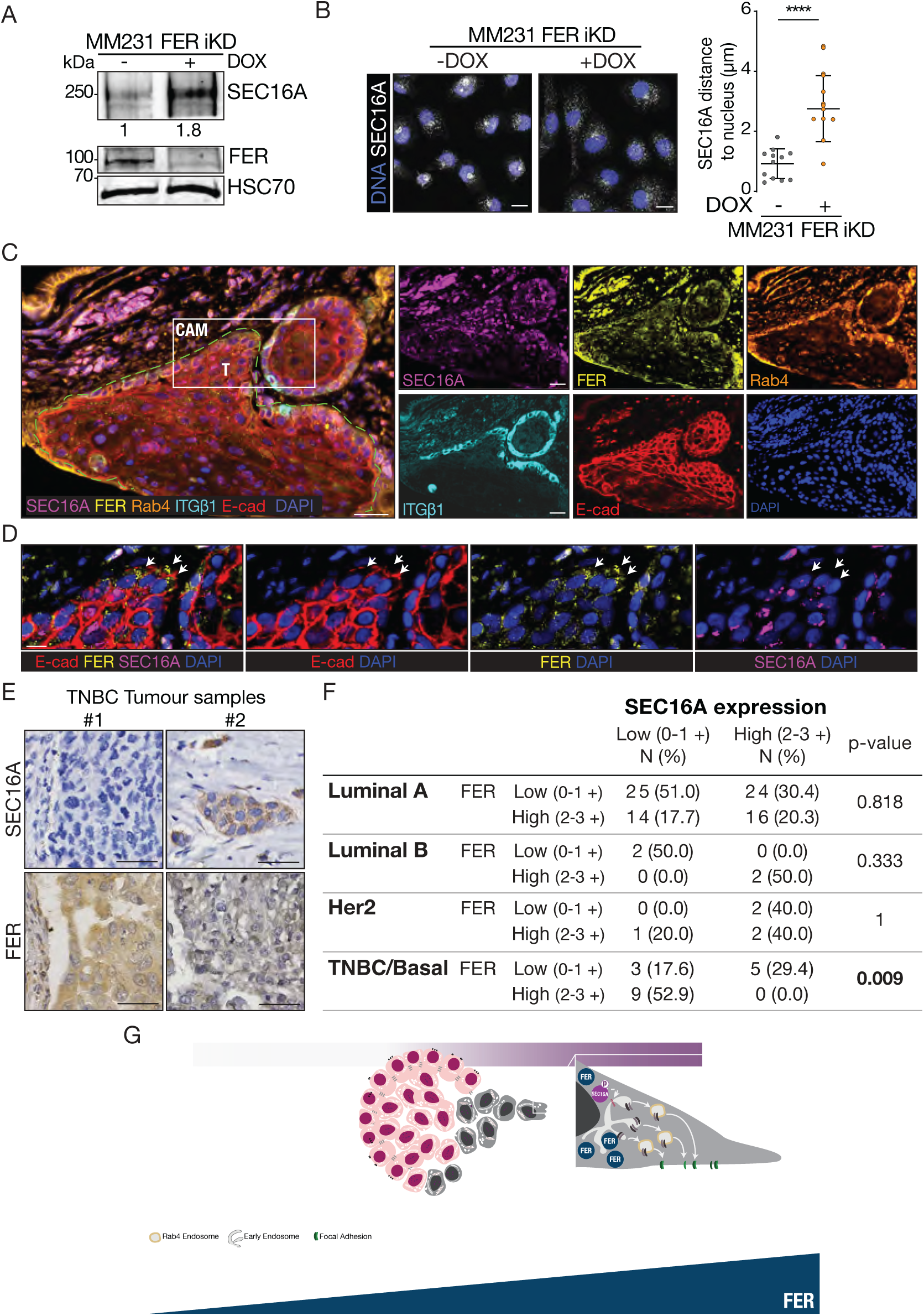
FER and SEC16A levels are inversely correlated in invasive TNBC cells. **(A)** Western blot on protein extracts from MM231 FER-iKD cells untreated or treated with DOX, blotted with anti-SEC16A or anti-HSC70 used as a loading control. Numbers on the bottom indicate the fold-changes in SEC16A levels normalised to HSC70 levels for each experimental condition. **(B)** (Left panels) Confocal images of MM231 FER-iKD cells untreated or treated with DOX, stained with DAPI (blue) and anti-SEC16A (grey). Scale bars represent 15 µm. (Right panel) Quantification of the shortest distance between SEC16A signals and the nucleus in MM231 FER-iKD cells treated with DOX (n=113) or without DOX (n=188). Data are from three biological replicates and include at least three fields analysed per condition and per replicate. Each dot represents one field. Error bars indicate standard deviation. Statistical significance was calculated using unpaired t-test. ****p<0.0001. **(C and D)** Multispectral immunofluorescence imaging of a TNBC PDXO #1 tumour xenograft invading on CAM. Scale bars represent 30 µm. Tissue samples were stained with anti-SEC16A (magenta), anti-FER (yellow), anti-Rab4 (orange), anti-Integrin β1 (cyan blue), anti-E-cad (red) and DAPI (blue). **(D)** Inset images highlight the levels of FER and SEC16A in leader invading cells. White arrows indicate leader cells. Scale bars represent 10 µm. **(E)** IHC images of TNBC clinical specimens, stained for SEC16A and FER. Scale bars: 50 µm. **(F)** Correlation between SEC16A and FER expression in Luminal A, Luminal B, HER2 and TNBC/basal breast cancer. Statistical significance estimated by **χ**2-test. P-values are indicated. Statistically significant p-values are indicated in bold. **(F)** Model by which phosphorylated SEC16A acts, downstream of FER, to coordinate the formation of Rab4-positive endosomes and focal adhesions and sustain invasion in TNBC. In TNBC, high levels of FER correlate with low SEC16A levels. Still, a certain amount of SEC16A protein is necessary to sustain FER-dependent endosomal recycling and invasion of integrins in TNBC.

We then asked if the inverse correlation between FER and SEC16A levels could also be observed in a series of breast cancer tissues. Strikingly, while FER and SEC16A levels were not inversely correlated in Luminal A, Luminal B and HER2 breast cancer subtypes, FER and SEC16A levels were significantly correlated in the TNBC/Basal subtypes (Fig 6E and F). These observations suggest that FER regulates SEC16A levels and subcellular distribution in TNBC.

## Discussion

Here we used clinically relevant TNBC PDXO models that preserve the heterogeneity and tumour architecture observed in patients^38^ to show that membrane trafficking is mainly regulated through phosphorylation in invasive FER-expressing TNBC PDXO cells and to identify SEC16A as a key involved player. Cluster 8 from the basal cancer cell population emerged as the likely pro-invasive population within TNBC PDXOs. This cluster upregulates the expression of ECM-related genes (FN1, CYP1B1 and TIMP1), and mesenchymal genes (ZEB1, VIM and KRT14), while downregulates the expression of differentiation (CD24) and epithelial (CDH1) genes. Consistent with our observations, FN1 and VIM are upregulated in invasive cells from MMTV-PyMT tumour organoids cultured in collagen-I, compared to those in Matrigel by scRNA-seq.^25^ Furthermore, CYP1B1 is expressed in pro-invasive TNBC cell lines, where it enhances their mesenchymal phenotype.^39,40^ Yet, pathways associated with membrane trafficking and cellular organisation are not recovered in any of the 8 basal cancer cells clusters identified by scRNA-seq. Also, these pathways are not regulated at transcriptional levels in cluster 8 between invasive and non-invasive PDXOs. Still, higher RAB4A levels could be required for cell invasion, as RAB4A is upregulated in cluster 8 from invasive TNBC PDXOs. RAB4A upregulation aligns with our subcellular structural analysis, which shows a striking enrichment of Rab4-positive recycling tubules in PDXO cells in contact with Collagen-I grown in invasive condition. Moreover, Rab4-positive endosomes have been proposed to serve as intracellular hubs to mediate the fast adaptation of TNBC invasive cells to extra-cellular cues.^41^ So, the upregulation of RAB4A in invasive FER-expressing TNBC PDXOs may exacerbate the recycling of integrins, thereby sustaining cell invasion and facilitating tumor spread. Beyond RAB4A upregulation, the invasive behaviour of TNBC PDXOs is also associated with significant phosphorylation/dephosphorylation events in proteins regulating pathways involved in the biogenesis and organization of cellular structures. These post-translational modifications appear to occur independently of changes in total protein abundance, as our proteomic analysis does not reveal a corresponding signature related to the biogenesis or organization of cellular structures. Instead, invasive PDXOs accumulate proteins associated with various metabolic processes, likely reflecting adaptation to nutrient-rich culture conditions. Of note, although western blot analysis indicates that FER accumulates in invasive PDXOs, FER is not detected in the full proteome dataset, possibly due to the limited detection power for proteins of low abundance by MS. This preferential regulation of ESR pathways through phosphorylation in FER-expressing TNBC PDXOs, rather than transcriptional changes, likely reflects a fast mechanistic adaptive response to environmental cues. Phosphorylation, as a reversible and rapid modification, allows cells to quickly adjust to changes in their surroundings, such as growth factor availability, or mechanical forces, without requiring the longer timescales associated with transcriptional reprogramming.^42^ Such fast mechanistic adaptation would be well-suitable for invasion, enabling accurate delivery and dynamic turnover of adhesion molecules at the plasma membrane. Importantly, invasive PDXOs are enriched in phosphorylation events on proteins involved in vesicle endocytosis (e.g. AAK1 and GAK, FCHO2, BIN1, among others), an initial step in receptor recycling and integrin turnover.^2^ In addition, the phosphorylation of CSNK1D and TRAPPC4, two proteins associated with COPII vesicle coating^43,44^, could play a pivotal role in TNBC invasion. CSNK1D, together with R-spondin 3 present in our pro-invasive media, could drive TNBC cell invasion through Wnt/β-catenin signalling, as both are potent inducers of Wnt/β-catenin signalling promoting invasion in TNBC cells.^45^ TRAPPC4 phosphorylation may be another mechanism controlling proteins recycling in invading TNBC cells. In colorectal cancer, TRAPPC4 acts as a scaffold between cargo proteins (e.g., PD-L1) and Rab11 in recycling endosomes^46^ to the plasma membrane to ensure proper surface expression of proteins. In addition, the hypo-phosphorylation of proteins associated with vesicle budding from the membrane in invasive PDXOs could regulate the trafficking of vesicles carrying matrix metalloproteinases that remodel the ECM, integrins, and growth factors. The fast response of TNBC cells to changes in the microenvironment might also be mediated by proteins bearing SH3-binding domains, which have been associated with endocytosis and TNBC chemoresistance^47,48^, and by Myosin II binding proteins, which can phosphorylate known tumour suppressors maintaining cell polarity and controlling epithelial-mesenchymal transition.^49,50^ Together, our findings suggest that the differential phosphorylation of the membrane trafficking machinery supports the invasive behaviour of FER-expressing TNBC cells.

Notably, our findings are in agreement with a model by which the ER export factor SEC16A functions as a FER effector driving TNBC cell invasion. Alike FER^7,9,35^, SEC16A promotes cell invasion and enhances the number of Rab4-positive endosomal tubules in MM231 cells. SEC16A is also required for the invasion of FER-expressing TNBC PDXO cells. Thus, SEC16A could promote cell invasion by controlling the spatial distribution of α6/β1-integrins downstream of FER. Consistent with this hypothesis, SEC16A has been proposed to stimulate α-integrin exocytosis in *Drosophila*.^51^ SEC16A is also known for its roles in protein transport from the ER to the Golgi, in mediating COPII vesicle formation at the transitional ER^52^ and in promoting protein trafficking from the perinuclear recycling endosome/trans-Golgi compartment back to the plasma membrane.^53^ Some of these functions, but not all, could depend on SEC16A interaction with SEC13.^54,55^ SEC16A activity is also controlled through phosphorylation on diverse Serine and Threonine.^52^ We propose that SEC16A is hyperphosphorylated downstream of FER, leading to integrin recycling and subsequent invasion. In invasive TNBC cells, the hyperphosphorylation of SEC16A could generate peptide’s negative charge and hydrophilicity hindering the identification of SEC16A S1327 phosphorylation by MS. Consistent with this model, MS could detect S1327 phosphorylation in FER-depleted MM231^9^ and in non-invasive TNBC PDXOs. Moreover, expressing a S1327 phospho-deficient version of SEC16A reduces the number of FAs and Rab4-positive endosomal tubules. Yet, the precise mechanism by which FER-dependent SEC16A phosphorylation controls the balance between endocytosis and recycling to regulate the amount of integrins on the cell surface remains to be determined. In addition to control SEC16A phosphorylation, FER keeps in check SEC16A protein levels and facilitates SEC16A perinuclear localisation in MM231 cells. Consistent with these observations, FER and SEC16A levels are inversely correlated in outer cells from TNBC PDXOs xenografted on CAMs and in TNBC samples specifically. Since depletion of either FER^7,9^ or SEC16 impairs proliferation, invasion, and affects both focal adhesions and Rab4-positive endosomal tubules, SEC16A accumulation outside the perinuclear region in cells with low FER levels may be non-functional. Yet we cannot exclude that this non-perinuclear SEC16A pool is involved in additional processes.

From a therapeutic perspective, SEC16A represents a promising target to treat a subgroup of TNBC. FER and SEC16A levels could distinguish one subgroup of highly invasive TNBC comprising one third of all TNBC cases. TNBC expressing high FER levels show better outcome after adjuvant taxane-based combination chemotherapy.^7,9^ Yet, these therapies display undesirable and long-term side effects, impacting the patient’s quality of life. In contrast, the small molecule Retro-2, which acts as a selective inhibitor of SEC16A^55^ shows no toxicity in either organs or blood from healthy mice, while strikingly abrogates the growth of TNBC cells in multiple preclinical models.^54^ Thus, Retro-2 could represent a selective and less toxic therapeutic strategy to inhibit the growth and invasion in FER-expressing TNBC.

In summary, this study highlights the key contribution of the post-translational regulation of membrane trafficking in sustaining the invasion of FER-expressing TNBC and features SEC16A as a promising TNBC therapeutic target.

## Supporting information

Supplemental figures with legends

## ACKNOWLEDGMENTS

This work was supported by funds from Fundação para a Ciência e Tecnologia (FCT) (grant number: COMPETE2030-FEDER-00667100 and EULAC/0002/2022), by funds from the Portuguese Medical Association and the BIAL Foundation (Maria de Sousa Award); Gilead Genese (grant number 17773); and the L’Oréal Portugal Medals of Honor for Women in Science to ST; and by funds from the Dutch Cancer Society (KWF 2020-13552) to AAK. ST was the recipient of 2020.02149.CEECIND, from FCT. FJ was the recipient of 2022.06763.CEECIND, from FCT. AS was the recipient of a fellowship from FCT 2025.02265.BDANA.

The authors acknowledge the support of the i3S Translational Cytometry (TraCy) Scientific Platform, the i3S Scientific Platform HEMS and i3S Scientific Platform Advanced Light Microscopy, member of the national infrastructure PPBI-Portuguese Platform of BioImaging (supported by POCI-01-0145-FEDER-022122). This work is a result of the GenomePT project (POCI-01-0145-FEDER-022184), supported by COMPETE 2020 - Operational Programme for Competitiveness and Internationalization (POCI), Lisboa Portugal Regional Operational Programme (Lisboa2020), Algarve Portugal Regional Operational Programme (CRESC Algarve2020), under the PORTUGAL 2020 Partnership Agreement, through the European Regional Development Fund (ERDF), and by Fundação para a Ciência e a Tecnologia (FCT). The scRNA-sequencing was performed at the Genomics i3S Scientific Platform with the assistance of Ana Mafalda Rocha. The transmission electron microscopy was performed at the HEMS core facility at i3S, University of Porto, Portugal with the assistance of Sofia Pacheco e Rui Fernandes.

The breast tissue and data bank at the Goodman Cancer Institute-Research Institute of McGill University Health Cantre (MUHC) is supported by the Database and Tissue Bank Axis of the Réseau de Recherche en Cancer of the Fonds de Recherche du Québec-Santé and the Québec Breast Cancer Foundation and certified by the Canadian Tumor Repository Network (CTRNet).

## Author contributions

Conceptualization, S.T..; methodology, S.T.; investigation, S.T. C.E., G.M., J.C., A.S., T.A.N., S.L., P.H., R.M.v.E., A.R.A., A.M., P.H. S.v.K., K.v.d.G., R.F., A. J. M. R.; formal analysis, S.T., B.C., B.S.M.; resources, P.v.D., A.A.K., M. P. and AMF; writing—original draft, S.T. and FJ; writing—review and editing, S.T., P.W.B.D. and F. J.; funding acquisition, ST; supervision, S. T.

