## Supplemental figures with legends for "Multi-omics analysis of TNBC organoids identifies phosphorylation of the membrane trafficking machinery as key event associated with FER-mediated invasion"

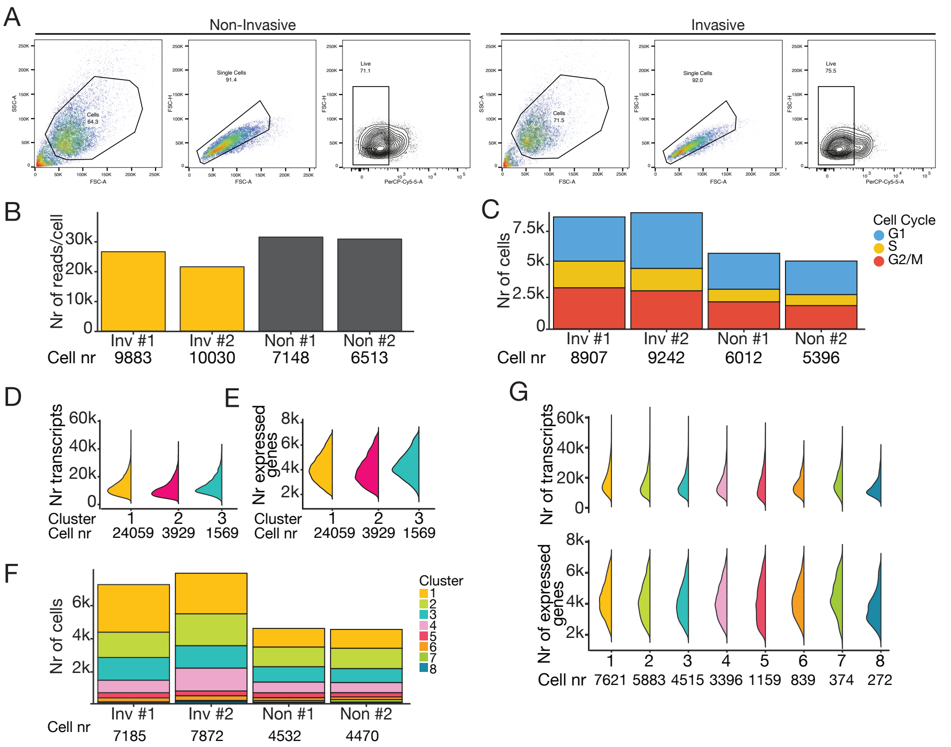


**Supplementary Figure 1: scRNA-seq data quality (A)** Flow cytometry gating strategy to isolate cells for scRNA-seq. **(B)** Plot of the total number of reads per cell for two PDXOs #1 replicates cultured in invasive (Inv #1 and Inv #2) and non-invasive (Non #1 and Non # 2) conditions. The number of cells analysed for each sample is indicated below the plot **(C)** Plot of the number of cells in each phase of the cell cycle for two PDXOs #1 replicates cultured in invasive (Inv #1 and Inv #2) and non-invasive (Non #1 and Non # 2) conditions. **(D and E)** Plots depicting the distribution of the number of transcripts counts **(D)**, or the number of expressed genes **(E)** in cluster 1 (Basal cancer cells), 2 (Luminal cells) and 3 (Myoepithelial cells)**. (F)** Plot of the number cells within each cluster belonging to the basal cancer cell population for two PDXOs #1 replicates cultured in invasive (Inv #1 and Inv #2) and non-invasive (Non #1 and Non # 2) conditions. **(G)** Plots depicting the distribution of the number of transcripts counts (upper panel) and the number of expressed genes (lower panel) for the 8 clusters constituting the basal-cancer cell PDXO populations. The number of cells for each cluster is indicated on the bottom. Related to Methods Sections “Single-cell isolation for mRNA-seq and flow cytometry analysis” and “scRNA-seq data analysis”.

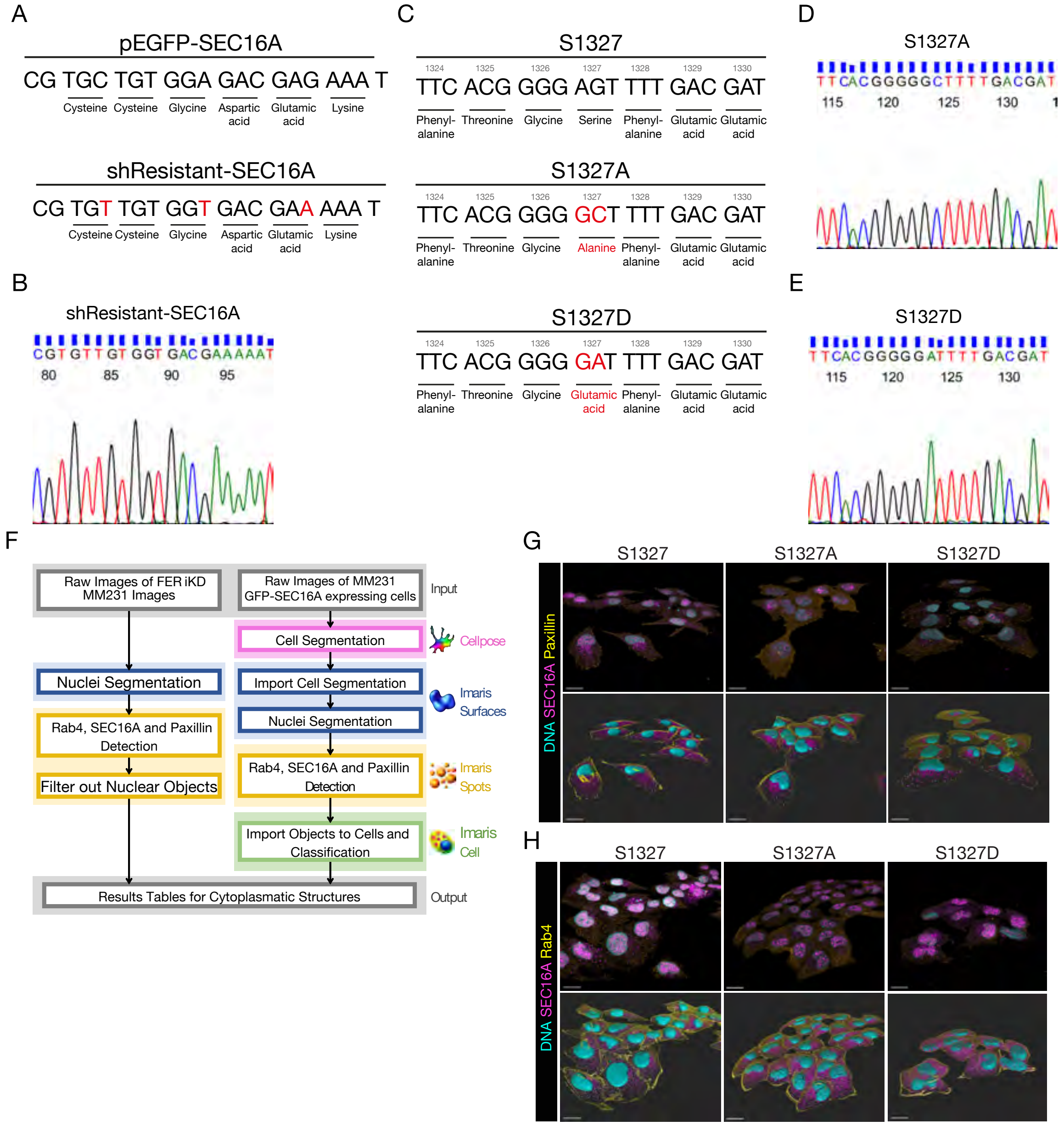


**Supplementary Figure 2 - Site-directed mutagenesis of SEC16A phospho-mutants and spinning disk confocal image data quantification strategy. (A)** Wild-type forward sequence of GFP-SEC16A and the corresponding shResistant-SEC16A construct generated by introducing synonymous mutations (mutated nucleotides highlighted in red). **(B)** Electropherogram of shResistant-SEC16A colony following site-directed mutagenesis, displaying the forward sequencing results. **(C)** Wild-type forward sequence surrounding the S1327 in EGFP-SEC16A and the engineered phospho-deficient (S1327A) and phospho-mimetic (S1327D) mutants (mutated nucleotides highlighted in red). **(D and E)** Electropherograms showing forward sequencing results of individual colonies for the S1327A (D) and S1327D (E) mutants after site-directed mutagenesis. **(F)** Graphical summary of the pipelines used to quantify Rab4, SEC16A and Paxillin signals from spinning disk confocal image data of MM231 and FER-iKD cells and GFP-SEC16A expressing cells. For MM231 cells, nuclei were segmented in Imaris, followed by detection of Rab4, SEC16A, and basal Paxillin puncta, with nuclear objects excluded from analysis. In FER-iKD cells, whole-cell segmentation was performed using Cellpose-SAM and imported into Imaris, followed by nuclear segmentation and puncta detection. Detected structures were assigned to individual cells for single-cell quantification of cytoplasmic features. After object import, cells were classified prior to quantification. **(G and H)** Confocal spinning-disk images of MM231 cells. **(G)** Upper panels show volume renderings of raw multichannel images with DAPI (cyan), SEC16A (magenta), and Paxillin (yellow). Lower panels display Imaris segmentation outputs with cell surfaces (outlined in yellow), nuclei (cyan), and SEC16A puncta (magenta) and Paxillin (yellow). **(H)** Upper panels show volume renderings of raw multichannel images with DAPI (cyan), SEC16A (magenta), and Rab4 (yellow). Lower panels display the corresponding Imaris segmentations with cell surfaces (yellow), nuclei (cyan), and puncta for SEC16A (magenta) and Rab4 (yellow). Variants shown are GFP-SEC16A-WT (WT), GFP-SEC16A-S1327A (S1327A) and GFP-SEC16A-S1327D (S1327A). All images were prepared in Imaris, with linear adjustments to brightness and contrast applied equally across conditions for presentation. Scale bars, 20 µm.

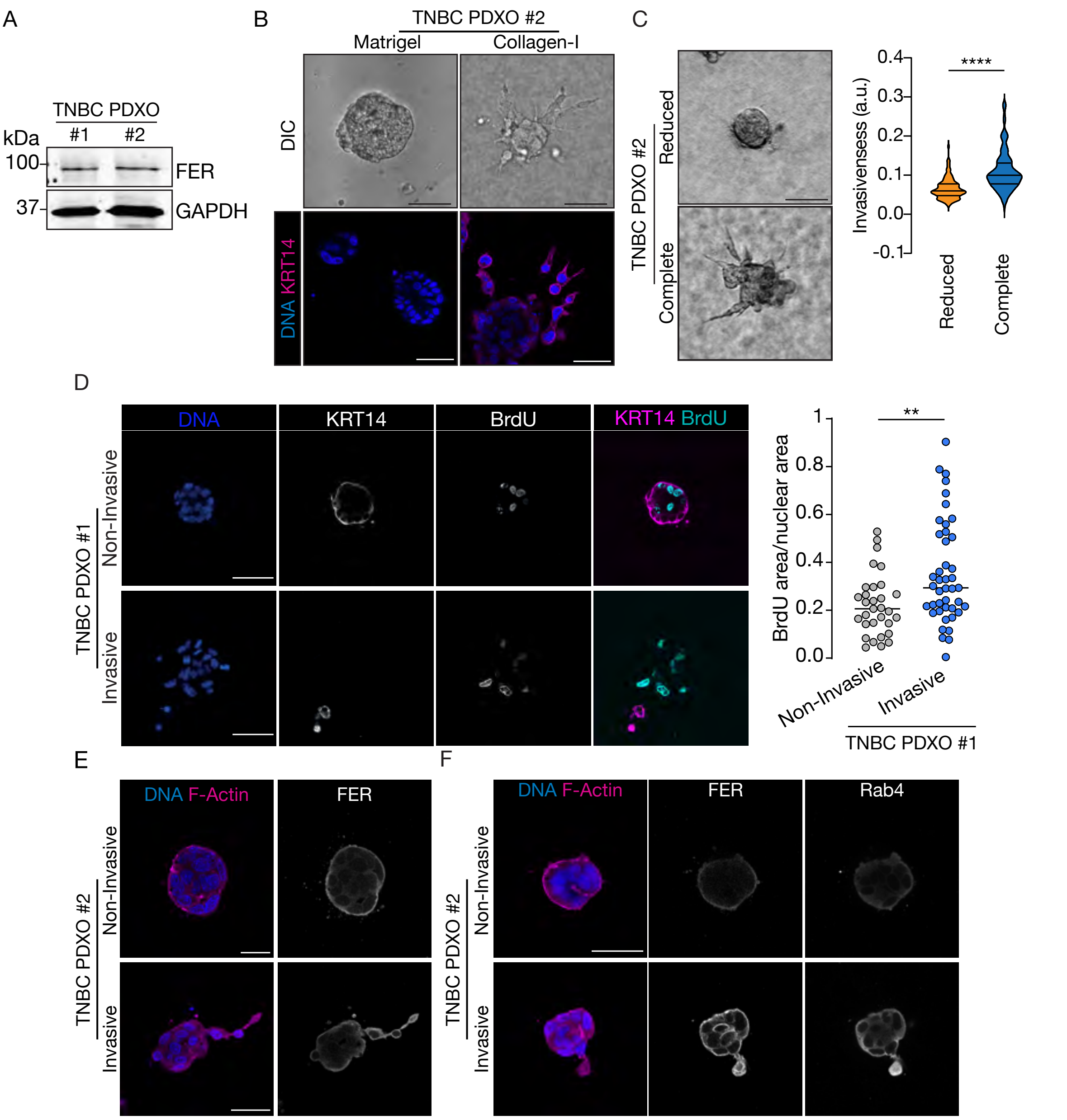


**Supplementary Figure 3: FER-expressing TNBC PDXOs invade in Collagen-I when cultured in Complete medium and display increased proliferative abilities. (A)** Western blot on protein extracts from TNBC PDXOs #1 and #2, blotted with anti-FER and anti-GAPDH used as loading control. **(B)** (Upper panels) DIC or (lower panels) confocal images of PDXOs #2 cultured in Complete medium and Collagen-I or Matrigel. (Lower panels) PDXOs #2 are stained with DAPI (blue) to stain the nuclei and anti-KRT14 (magenta) to identify basal-like cells. Scale bars represent 50 µm. **(C)** (Left panels) DIC images of PDXOs #2 cultured in Reduced or Complete medium and Collagen-I. (Right panel) Quantification from three biological replicates of the invasiveness of PDXO #2 cultured in Reduced or Complete media and Collagen-I. At least 110 PDXOs were quantified. *p<0.0001. Statistical significance was calculated using Mann-Whitney test. **(D)** (Left panels) Confocal images of PDXOs #1 cultured in Reduced or Complete medium and Collagen-I, stained with DAPI (blue), anti-BrdU (grey or cyan) and anti-KRT14 (grey or magenta). Scale bars represent 50 μm. (Right panel) Quantification of the number of BrdU positive nuclei in PDXOs #1 cultured in Reduced (n=31) or Complete (n=43) medium and Collagen-I. Statistical significance was calculated using unpaired t-test. **p=0.0031. **(E)** Confocal images of PDXOs #2 cultured in Reduced or Complete medium and Collagen-I, stained with DAPI (blue), Phalloidin (magenta) and anti-FER (white). **(F)** Confocal images of PDXOs #2 cultured in Reduced or Complete medium and Collagen-I, stained with DAPI (blue), Phalloidin (magenta), anti-FER (white) and anti-Rab4 (white). Scale bars represent 50μm. Related to Figure 1.

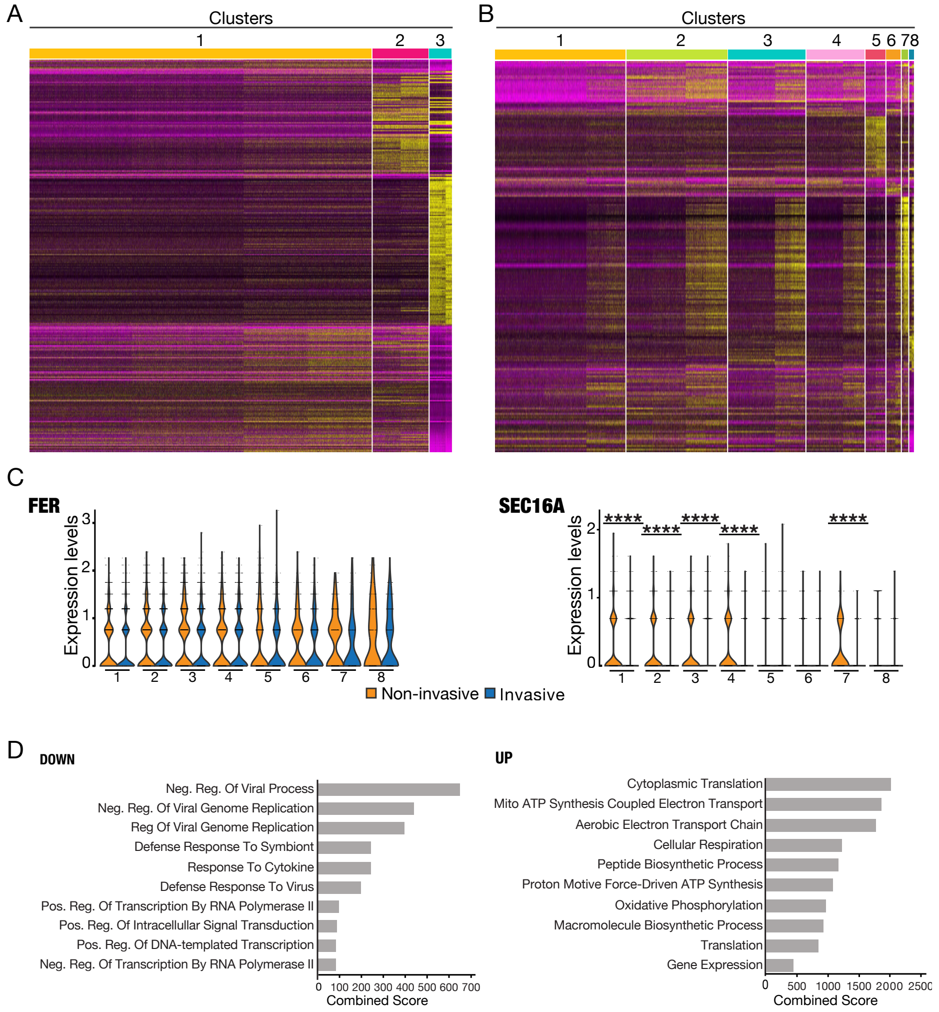


**Supplementary Figure 4: scRNA-seq clusters characterization (A)** Heatmap of scaled gene expression data. Plotted are all differentially expressed genes between the three main clusters (1–Myoepithelial cells (yellow); 2-Luminal cells (magenta); 3-Basal Cancer Cells (cyan) in PDXOs #1 cultured in invasive and non-invasive conditions. **(B)** Heatmap of scaled gene expression data. Plotted are all differentially expressed genes between the 8 clusters within the basal cancer cells compartment. **(C)** Quantification of FER and SEC16A expression between the 8 clusters of the basal cancer cells compartment of PDXOs cultured in invasive (blue) and non-invasive (yellow) conditions. The y axis represents the distribution of expression through a density plot. Differential expression analysis between cells cultured in invasive and non-invasive conditions was carried out using the FindMarkers Seurat function for each identified cluster. ****$P < 0.0001$. **(D)** Gene enrichment analysis of up- or down-regulated genes in cluster 8 between invasive and non-invasive PDXOs. Significantly altered pathways were filtered for adjusted p-value (<0.05). Related to Figure 3.

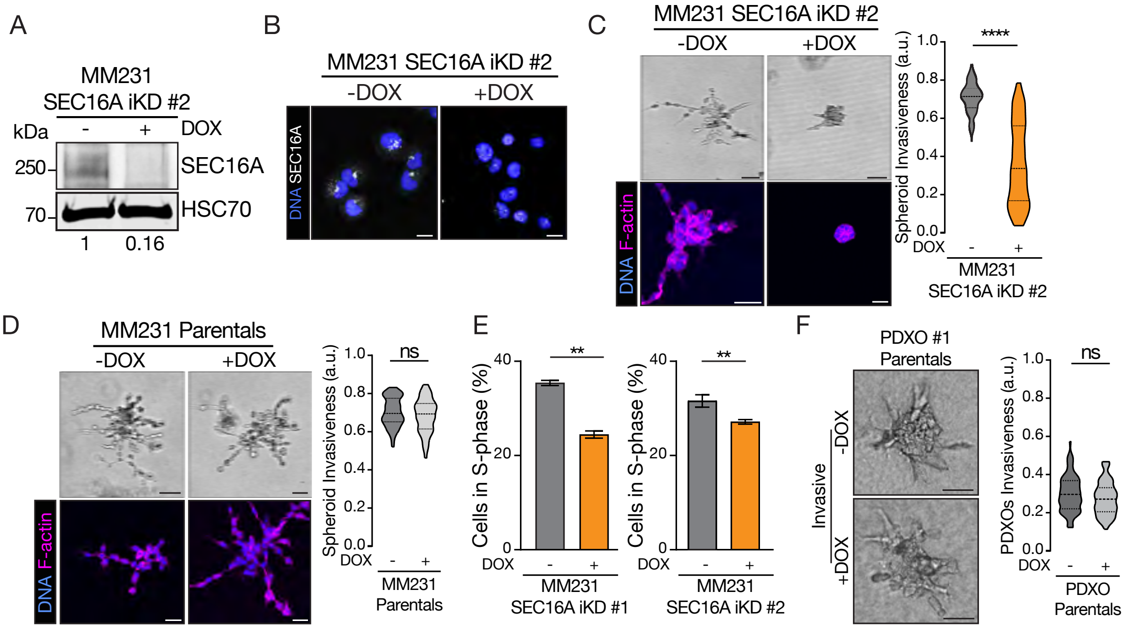


**Supplementary Figure 5: SEC16A depletion reduces cell invasion and proliferation in TNBC cells. (A)** Western blot on protein extracts from MM231 SEC16A-iKD #2 cells untreated or treated with DOX, blotted with anti-SEC16A and anti-HSC70 used as a loading control. Numbers below indicate the fold-changes in SEC16A levels for each condition. **(B)** Confocal images of MM231 SEC16A-iKD #2 cells untreated or treated with DOX, stained with DAPI (blue) and anti-SEC16A (white). Scale bars indicate 15 μm. **(C)** (upper left panels) DIC or (lower left panels) confocal images of MM231 SEC16A-iKD #2 spheroids in Matrigel untreated or treated with DOX. (Lower left panels) spheroids are stained with DAPI (blue) and Phalloidin (magenta). Scale bars indicate 50 μm. (Right panel) Quantification of the invasiveness of MM231 SEC16A-iKD #2 spheroids in Matrigel untreated or treated with DOX. Data are from three biological replicates. Statistical significance was calculated using unpaired t-test. ****$p<0.0001$. **(D)** (Upper left panels) DIC or (lower left panels) confocal images of MM231 parental spheroids in Matrigel untreated or treated with DOX. (Lower left panels) Spheroids are stained with DAPI (blue) and Phalloidin (magenta). Scale bars indicate 50 μm. (Right panel) Quantification of the invasiveness of parental MM231 spheroids in Matrigel untreated or treated with DOX. Data are from three biological replicates. Statistical significance was calculated using unpaired t-test. ns indicates non-significant ($p=0.2219$). **(E)** Quantification of the number of cells in S-phase in MM231 SEC16A-iKD #1 and MM231 SEC16A-iKD #2 cells untreated or treated with DOX. Data are from three biological replicates. Statistical significance was calculated using unpaired t-test. **$p=0.0038$ between untreated and DOX-treated MM231 SEC16A-iKD #1. **$p=0.0053$ between untreated and DOX-treated MM231 SEC16A-iKD #2. **(F)** (Left panels) DIC images of parental TNBC PDXOs #1 untreated or treated with DOX, cultured in invasive condition. (Right panel) Quantification of the invasiveness of parental PDXOs untreated or treated with DOX**.** Data are from three biological replicates with more than 30 PDXOs analysed per condition and per replicate. Statistical significance was calculated using unpaired t-test. ns indicates non-significant (p= 0.0724). Related to Figure 4.

### Fig 1G

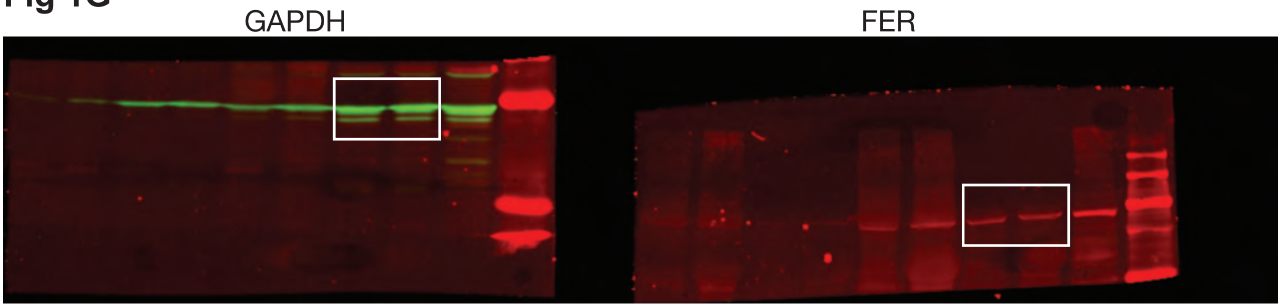


### Fig 3A

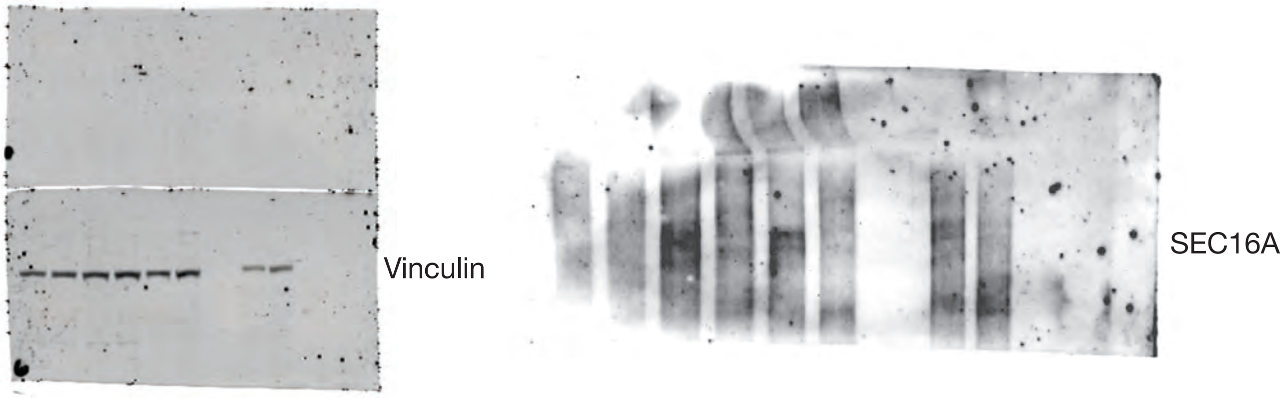


### Fig 3I

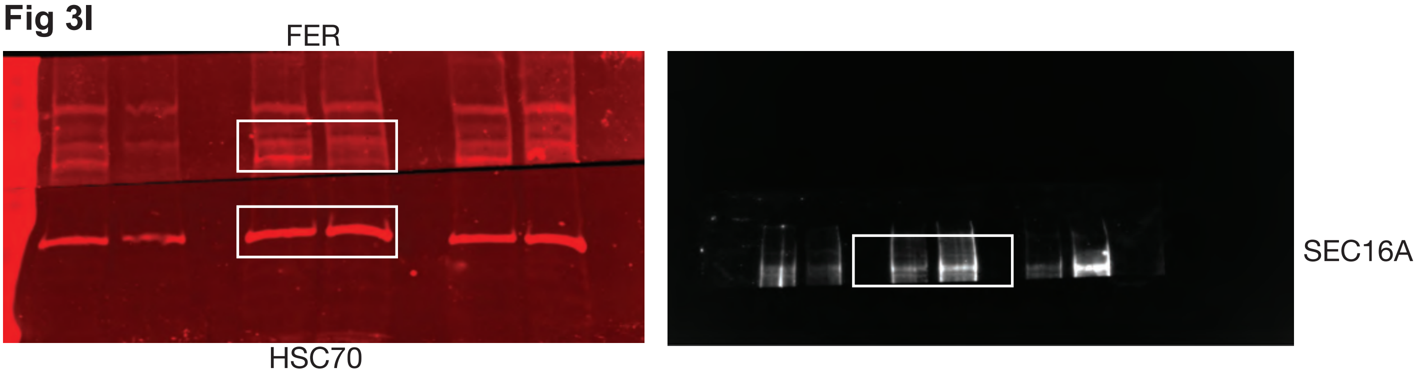


### Fig 3L

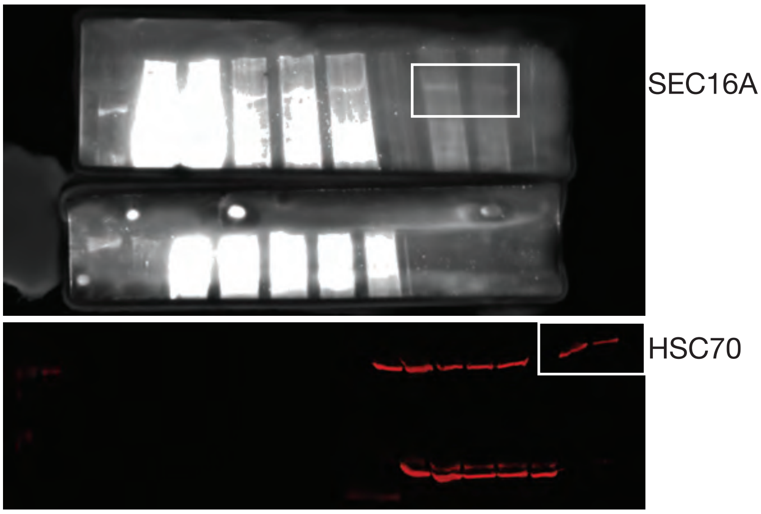


### Fig S3A

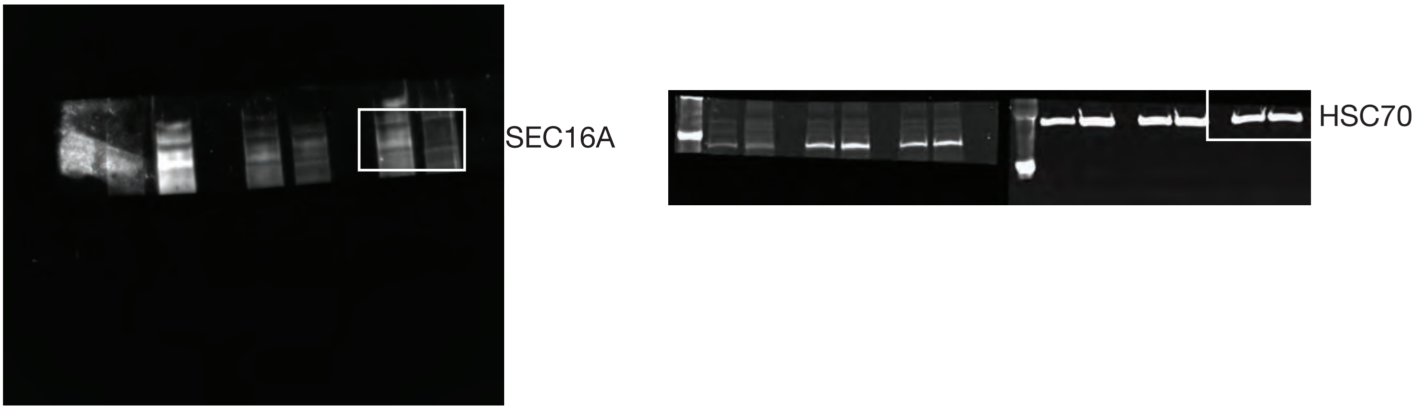
